# Isolation of Extracellular Vesicles from Minimal Volume Ascites Fluid Using Strong Anion Exchange Beads

**DOI:** 10.1101/2025.09.24.678291

**Authors:** Tyler T. Cooper, Lorena Veliz, Farzaneh Afzali, Cédric Djuomessi, Owen F.J. Hovey, Robert Myette, Tiffany P.A. Johnston, Chris Wells, Tristan Robertson, Dylan Burger, Sheela A. Abraham, Trevor G. Shepherd, Andrew Craig, François Lagugné-Labarthet, Gilles A. Lajoie, Lynne-Marie Postovit

## Abstract

Ovarian cancer (OC) remains a leading cause of gynecologic cancer mortality due to late-stage diagnosis and limited early detection strategies. Ascites fluid, a pathological hallmark of OC, is a rich source of tumor-derived extracellular vesicles (EVs) that reflect the tumor microenvironment and hold promise for biomarker discovery. However, isolating EVs from minimal ascites volumes (<100 µL) poses technical challenges using conventional methods like ultracentrifugation or size-exclusion chromatography (SEC). This study explores the application of strong anion exchange (SAX) magnetic beads (Mag-Net) for efficient EV isolation from as little as 2 µL of ascites fluid from both murine models and a human patient with mucinous borderline tumor. We demonstrate that SAX achieves robust EV capture at 10µl of input volume, enabling comprehensive proteomic profiling and single-EV surface-enhanced Raman spectroscopy (SERS) with a >2-fold increase in proteomic depth compared to raw ascites. Notably, this study was able to identify 1000 proteins not previously annotated in Vesiclepedia for OC-derived EVs, alongside distinct SERS signatures, highlighting the potential for multiomic analysis. Comparative analysis with UC revealed enhanced proteomic depth obtained with SAX beads, albeit we also observed differential detection of canonical markers (e.g., CD9, CD81) between input volumes of ascites fluid. These findings establish SAX as a scalable, low-input platform for EV-based biomarker discovery, paving the way for improved early detection and molecular insights into OC progression.

## 1, Introduction

Ovarian cancer (OC) remains one of the most lethal gynecologic malignancies worldwide, accounting for 4.7% of cancer related deaths in 2020^1^. >90% of OC is epithelial in origin, with high-grade serous carcinoma (HGSC) the most aggressive and prevalent cell subtype. Fallopian tubes are the purported site of origin for HGSC; however, endometrial and non-ovarian tissues contribute to the remaining OC histotypes of distinct etiology^2^. Frequent diagnosis of aggressive OC at advanced stages contributes to poor survival rates, largely due to the absence of effective early detection strategies and reliable biomarkers^3^. This underscores an urgent need for innovative methodologies to identify biomarkers that can enable earlier diagnosis in order to improve patient outcomes. Ascites fluid is a common pathological feature of OC and accumulates in the peritoneal cavity where it serves as a rich biological reservoir of tumor-derived material, including extracellular vesicles (EVs), cytokines, and metabolites^4-7^. The molecular composition of ascites fluid is largely reflective of a malignant vs benign tumor microenvironment, making ascites an ideal substrate for studying disease progression, defining histological subtypes, and uncovering potential diagnostic markers. An increased proximity of isolated EVs relative to the tumor is particularly valuable for early biomarker discovery since tumor-specific or tumor-associated antigens may efflux from tumor microenvironment into systemic circulation^8^. To this end, our recent publication utilized the proteomes of cell culture and primary ascites EVs to develop targeted proteomics workflows that identified combinatorial biomarkers, such as Mucin-1 (MUC1), on plasma-derived EVs from donors with early-stage HGSC^8^. Likewise, we have developed workflows to assess single-EVs using surface enhanced Raman spectroscopy (SERS) with the goal of increasing the sensitivity of detecting early-stage disease^9, 10^. Additional “omic” analysis by others have uncovered nucleic acids and lipid signatures distinct to early-stage disease^11, 12^. Taken together, EVs represent an attractive source of biomarkers which requires adopting improved methodologies to increase EV purity.

EVs are nanoscale particles secreted by nearly all cell types and have become increasingly recognized as key players in cancer biology due to their role in intercellular communication^13^. EVs carry diverse bioactive cargo, such as proteins, lipids, and nucleic acids, that mirrors the molecular state of their originating cells often carry unique signatures of disease. When isolated from ascites fluids from OC patients, EVs offer a snapshot of the tumor microenvironment that contains active sites of metastasis in mesothelial, adipose, or vascular/lymphatic systems^4, 14^. Dissemination of the primary tumor entails the aggregation of therapeutically resistant spheroids with a pseudo-senescent or dormant phenotype that reawaken upon fibroblast activation, adhesion to the mesothelium, or secondary sites of metasasis^6, 7, 15, 16^. Taken together, multiomic profiling of EVs isolated from biofluids (e.g., ascites, plasma, etc.) holds immense potential as a reservoir of molecular signatures that not only aid in early detection but also provide critical insights into mechanisms underlying disease progression and therapeutic resistance. In an ideal scenario, sampling early fluid accumulation in the peritoneal cavity or surrounding ovaries may uncover early indicators of OC^5, 6^. Unfortunately, isolating EVs from minimal volumes (<100µl) possess significant technical challenges, as traditional isolation methods (i.e. ultracentrifugation; UC, size-exclusion chromatography; SEC) may risk compromising EV cargo integrity and typically demand larger inputs for sufficient recovery. The most recent proteomic profiling of ascites fluid used >10mL to isolate EVs using UC or SEC^17^.

Utilizing the biophysical properties of EVs has been a focus to improve EV purity without compromising total yield. One example is asymmetric flow field-flow fractionation that utilizes the diffusion coefficients from two perpendicular liquid flows to separate and concentrate EVs^18^. However, the global adaptation of this approach is limited due to required technical expertise, cost prohibitions, and the need for sequential purification or concentration workflows^19^. Simplified approaches have targeted the lipid enclosed membrane of EVs using magnetic beads coated with hydrophilic and aromatic chemical motifs to improve the separation of EVs from lipoproteins, an often-co-isolated contaminant with similar biophysical properties^20^. One distinguishing characteristic of particles present in biofluids is the zeta-potential or overall negative surface charge of each particle class. EVs possess a more negative surface charge relative to HDL and LDL particles due to the presence of negatively charge lipid groups^21^. The enrichment of phosphatidylserine on the surface of cancer EVs^22^ and EVs released during periods of stress or apoptosis exacerbates this electrochemical delta^23^.Thus, exploiting this delta enriches for cancer-derived EVs in a robust and scalable manner. Wu *et al*. recently developed porous magnetic beads equipped with quaternary ammonium groups which interact with regions of negative surface charge using strong anion chemistry (SAX) at pH=6.0-6.5^24^. This workflow, termed Mag-Net, demonstrated a reproducible enrichment of EVs from plasma for the purposes of biomarker discovery in neurological disease. Others have employed Mag-Net workflows on alternative biofluids, such as urine and serum^25, 26^. To our knowledge, this is the first report of using SAX beads to isolate EVs from ascites fluid. Herein, this study exemplifies the application of SAX EV isolation and digestion on ascites samples from both a pre-clinical murine model and a human patient with borderline mucinous OC. Notably, we demonstrate the feasibility of EV capture and proteomics from as little as 2 μL of starting input volume and validate these workflows for alternative ‘omic’ analysis. Furthermore, as little as 10 μL of input volume supports single-EV SERS in parallel with comprehensive global proteomics^10^. Ultimately, this work provides foundational example using low input volumes to obtain mulitomic profiles of ascites EVs.

## 2. Experimentation

### 2.1 Murine Ascites

All animal experiments were conducted in accordance with protocols approved by the University Animal Care Committee at Queen’s University. A total of 10 mice were used across a survival cohort, 5 mice per condition. By intraperitoneal (IP) injections, 5 × 10^6^ ID8-Trp53^-^/^-^ dual-labeled tdTomato-Luc murine HGSC cells (shGFP or shPKR4653) suspended in 100µL PBS were madministered to 6–8-week-old female C57BL/6 mice (Charles River Laboratories, International Inc.). Body weight and abdominal girth were recorded twice weekly. Mice were euthanized when abdominal diameter reached ≥35 mm, indicative of substantial malignant ascites accumulation. Ascites fluid was collected either directly from the peritoneal cavity or after lavage with a known volume of PBS, using a 1mL syringe (BD 1ml TB Syringe; Cat# 309659) and 26-gauge needle (Precisionglide, Cat# 8024416). The collected fluid (typically 0.5–2 mL per mouse) was immediately transferred to a sterile 15 mL conical tube and kept on ice. To remove cells and debris, the ascites fluid was subjected to a series of centrifugation steps. First, samples were centrifuged at 300 × g for 10 minutes at 4°C to pellet intact cells. The supernatant was carefully transferred to a new tube and centrifuged at 2,000 × g for 15 minutes at 4°C to remove larger debris and apoptotic bodies. Finally, the supernatant was centrifuged at 10,000 × g for 20 minutes at 4°C to eliminate smaller debris and microvesicles, yielding a clarified ascites fluid enriched with EVs. The resulting supernatant was stored at -80°C for subsequent analysis. Samples were slowly thawed on ice and centrifuged at 10,000g for 5 mins to pellet debris before experimentation on supernatant.

### 2.2 Human Ascites

Ascites fluid was obtained from a consented adult donor diagnosed with a Stage1B mucinous borderline tumor. Patient consent for the use of clinical specimens in this study was obtained in accordance with to the Western University Research Ethics Board approved protocol (#115904). The patient underwent a laparotomy for the removal of ovarian masses, along with a hysterectomy, bilaterial salpingo-oophorectomy, and omentectomy, all performed by a trained gynecologic oncologist. Following the induction of general anesthesia, a vertical midline incision was made to access the peritoneal cavity. Using a suction apparatus connected to a sterile collection canister, approximately 50 mL of ascites fluid was collected. Ascites fluid (ranging from 10–50 mL per donor) was aspirated gently to avoid hemolysis or excessive shear stress that could compromise EV integrity, then immediately transferred to sterile 50 mL conical tubes and placed on ice. An additional three ascites and plasma samples, from donors with FIGO stage I high-grade serous carcinoma, were obtained from the gyneological biobank at CR-CHUM with approval from Research and Ethics Committee at CHUM (#2026-13303). To remove cells and debris, the fluid underwent sequential centrifugation. Initially, samples were centrifuged at 2000 × g for 10 minutes at 4°C to pellet viable cells and debris. The supernatant was carefully decanted into a fresh tube and centrifuged at 10,000 × g for 15 minutes at 4°C to eliminate larger debris and apoptotic bodies. The processed ascites fluid was either used immediately for EV isolation or aliquoted and stored at -80°C for subsequent experimentation.

### 2.3 Mag-Net

Strong anion exchange (SAX) beads were washed 6 times with Binding Buffer (BB;100mM bis-Tris-propane/150mM NaCl/pH=6.4). Processed ascites fluid (10-100µl) was diluted 1:1 with BB before incubation with beads in a 1:4 (bead/ascites) ratio, termed capture volume. Notably, the 1:4 is calculated based on the pre-diluted ascites fluid (i.e. 25µl beads in 100µl ascites fluid) for 10mins at 37°C on a thermomixer set at 600rpm. Unbound ascites fluid was collected and labelled as “Non-captured” or “NC” fraction throughout the study. Captured EVs were washed 4-6 times with Wash Buffer (50mM bis-Tris-propane/150mM NaCl). To limit EV shearing using WB, washing was performed off magnet once (2 minutes, 600rpm) in 2x capture volume three times followed by washing on-magnet in 2x capture volume. EVs were lysed on beads using SP3 protocol for proteomic analysis^27^, detailed below. For select experiments, EVs were eluted from beads using a 25mM bis-Tris-propane/ 1M NaCl/0.1% Tween-20 solution. A detailed protocol can be found here^28^.

### 2.4 Ultracentrifugation

For EV isolation by ultracentrifugation (UC), 100µl was diluted to 800µl with 1x PBS and centrifuged at 100,000 x g for 60 minutes using a MLA-130 rotor (130,000 rpm) for the OptimaMax benchtop ultracentrifuge (Beckman-Coulter) using 1mL polycarbonate tubes. Supernatants were carefully removed from EV pellets by pipetting the top 80% without disturbing the residual supernatant or EV pellet. The residual 20% of supernatant was pipetted from EV pellet by horizontal inversion of tube. EV pellets were collected with either 1x PBS, Mag-Net BB, or proteomic lysis buffer.

### 2.5 SP3 Proteolysis

Murine Ascites, NC, and EV fractions were lysed using a buffer comprised of 2% SDS/1% NP-40/150mM NaCl/pH=8.0. Human ascites, NC, and EV fractions were lysed in a buffer comprised of 4M Urea/2%SDS/1%NP-40/1%SDC/0.1%DDM/150mM NaCl/pH=8.0, unless otherwise specified. Protein precipitation and proteolysis were conducted according to Hughes *et al*. with slight modifications^27^. Protein precipitation was performed using either 70% ACN or 50% EtOH for 10 minutes on a thermoshaker (800 rpm) at 24 °C. Protein-bead isolated were washed on-magnet using 3 washes with 80% EtOH and a final wash of 95% ACN before the addition of digestion buffer comprised of 50mM Ammonium Bicarbonate/0.01% DDM, unless otherwise specified. Proteolysis was performed with LysC, 2 hours, 37 °C(900rpm) followed by Trypsin, 18 hours, 37 °C(900rpm). Digests were collected in fresh tubes and residual peptides were isolated from beads with water before pooling and acidification to 0.1% TFA prior to C18 stage-tip cleanup^8^.

### 2.6 Liquid Chromatography Coupled to Tandem Mass Spectrometry (LC-MS/MS)

The MS/MS spectra were obtained using the protocol previously reported by our group. Briefly, 10 µg of EV or 50 µg of cell lysates were isolated by single-pot protein precipitation (SP3) using 50% Ethanol and proteolysis (50 mM Ammonium Bicarbonate, 1:50 TrypLysC, 18 hours, 37°C) and analysed on a Thermo Eclipse using Gas-phase Fractionation operating in Data-Independent Acquisition (GPF-DIA), as previously described^29, 30^. Normalized HCD collision energy was set to 30. In select experiments, High field asymmetric ion mobility mass spectrometry (FAIMS) was operated in standard resolution with a gas flow of 4.6L/min and compensation voltage (CV) set to –50. Spectral libraries were first searched in DIA-NN against the human proteome before quantitative analysis using label-free quantification (MaxLFQ)^31^. The software parameters were set to a precursor mass tolerance of 20 ppm and a fragment ion mass tolerance of 10 ppm. The search parameters allowed for up to 2 missed cleavages and a maximum of 2 post-translational modifications per peptide. Missing value imputation, transformations, and proteomic data visualizations were performed using in-house Python code.

### 2.7 Surface Enhanced Raman Spectroscopy (SERS)

For the SERS measurements, the isolated EVs were diluted at 1:20 in ultrapurergwater, as previously described^10^. Then 10−20 μL of the solution were drop-casted onto the nanofabricated gold nanoholes array and dried for 30 min. The detailed fabrication of the Au nanoholes array in described is previous work ^10^. A confocal Raman microscope (XploRA Plus) was used to acquire the Raman spectra of each of the cell lines isolated by ultracentrifugation. An excitation laser with an excitation wavelength of 785 nm, a 600 groove/mm grating, and a 100×objective (N.A. = 0.9) were selected to perform the experiments. Additionally, a slit width of 100 μm and a pinhole of 300 μm were set to obtain the spectra with minimal background. The samples were raster scanned typically over a surface of (20 × 20) μm^2^ with an acquisition time of 4 s/spectrum, and a laser intensity of 5 × 10^5^ W/cm^2^. The scanning resolution was of 0.8 μm/spectrum along the X and Y direction yielding typically 360 spectra for a given map.

### 2.8 Nanoparticle Tracking Analysis (NTA)

NTA was performed on a ZetaView (Particle Metrix). In accordance with the manufacturer’s instructions, polystyrene 100 nm beads (Particle Metrix) were used for the daily calibration and instrument performance check. 5−20 μL of samples were diluted to 1 mL with 0.22 μm filtered PBS to obtain>400 particles per frame and measurements in the scatter mode were performed at RT at 11 positions. Triplicate cycles were acquired, and acquisitions with at least 9 out of 11 fields of view were retained for data analysis. Average hydrodynamic size and particle concentration were obtained by averaging technical triplicates for each donor pool.

### 2.9 Atomic Force Microscopy (AFM)

For the preparation of the samples, the isolated EVs were diluted at 1:20 in ultrapure water to reduce salt residue contamination when drying, as previously described^10^. Then, 10 μL of the solution was drop-casted into freshly cleaved mica substrate and let dry overnight. solution was drop-casted into a freshly cleaved mica substrate and let dry overnight. The AFM measurements were performed in a BioScope Catalyst atomic force microscope (Bruker). An NCLR-50 cantilever with a spring constant of 48 N/m and a resonance frequency of 190 kHz was selected for obtaining AFM images in tapping mode. The height images (topography) were obtained using a scan rate of 0.5 Hz and at 512 × 512 pixels. Finally, the images were processed by Gwyddion software. AFM was used for a qualitative analysis of EV-release from the surface of magnetic SAX beds and meant to complement our proteomic and NTA analyses.

### 2.10 Transmission Electron Microscopy (TEM)

20µl of EVs in elution buffer (100µl), minus 0.1% Tween-20, were cast dropped on to carbon grid (AGS160, Agar Scientific) and allowed to bind to surface during 10-minute incubation at room temperature. EV-bound grids were washed three times with 0.1M glycine in water for 5 minutes per washed, followed by five washed for 5 minutes each in MiliQ water. Carbon grids were counterstained using 2% urynal acetate in water for 2 minutes at room temperature before drying for at least 1 hour prior to imaging. Imaging was performed at McGill Facility for Electron Microscopy Research facility using FEI Tecnai G2 Spirit BioTwin 120 keV Cryo-TEM. Cell-derived, SEC-isolated EVs were kindly donated by Centre of Applied Nanomedicine at McGill for positive control of EV morphology under our staining protocol.

### 2.11 Scanning Electron Microscopy (SEM)

For SEM analysis, the magnetic beads containing EVs were diluted 1:20 in ultrapure water to reduce salt residue contamination when drying Later, 10 μL of the solution was deposited on a clean piece of silicon wafer and allowed to dry. After sample preparation, the Magnet beads and EVs were characterized using a LEO 1530 (Zeiss) scanning electron microscope with a 3.0 kV EHT voltage and a 30.0 μm aperture. SEM was used for a qualitative analysis of EV-binding to the surface of magnetic SAX beds and meant to complement our proteomic analyses.

### 2.12 Western Blot

A total of 15 μg of protein, determined using a BCA assay, was mixed with 5 μL of 6X Laemmli buffer, and the volume was adjusted to 30 μL. The samples were heated at 95°C for 10 minutes before being loaded onto a 12% SDS-PAGE gel (see Appendix A). The proteins were then transferred onto a polyvinylidene fluoride (PVDF) membrane for Western blotting, Amersham Hybond type, 0.45 μm (MilliporeSigma, cat. no. GE10600023), using a Trans-Blot Turbo semi-dry transfer system (Bio-Rad, cat. no. 1704150). The membranes were then incubated at 4°C for 15–16 h with primary antibodies against CD9 (Cell Signaling Technology, cat. no. 13403; dilution 1:1,000 in 5% bovine serum albumin [BSA], Wisent, cat. no. 800-095-EG, prepared in TBS-T) or albumin (Cell Signaling Technology, cat. no. 4929; dilution 1:1,000 in 5% non-fat milk prepared in TBS-T). TBS-T consisted of Tris-buffered saline containing 0.2% Tween-20. Following incubation with the primary antibodies, the membranes were washed four times with TBS-T for 10 minutes per wash. The membranes were then incubated with a horseradish peroxidase-conjugated anti-rabbit secondary antibody (Goat Anti-Rabbit IgG [H+L]-HRP, Jackson ImmunoResearch/Cedarlane, cat. no. 111-035-144). Detection was performed by chemiluminescence using ECL reagent (Pierce, 32209, ThermoFisher Scientific). Finally, the membranes were imaged using a ChemiDoc system (Bio-Rad).

### 2.13 Data Analysis

Raw LC-MS/MS files were converted to mzML files using msconvert with the following parameters: peakPicking:msLevel =1-, zeroSamples:removeExtra 1-. Downstream missing value imputation, enrichment analysis, and data visualization was performed in Python, using libraries including MissForest^32^ and GSEApy^33^.

## 3. Results and Discussion

### 3.1 SAX effectively enriches EVs from ascites fluid

This study evaluated the applicability of the Mag-Net protocol, utilizing SAX chemistry, to effectively isolate EVs from ascites fluid of both murine and human origin for multiomic analyses **(Figure 1A)**. Under slight acidity (pH=6.3-6.5) the capture of EV-like particles on the surface of SAX beads was observed using SEM **(Figure 1B)**; moreover, we occasionally observed a web-like binding of several beads mixed with surface bound particles **(Figure 1SA)**. It is essential that all studies involving EV-enrichment methodology perform EV characterizations according to the minimal guidelines established by the International Society of Extracellular Vesicles (ISEV)^34, 35^. Accordingly, ascites EV isolations were benchmarked against paired plasma samples to assess EV characteristics, such as size and cup-like morphology **(Figure 1C-D, Figure S1B-C)**. SAX-enriched EVs represent a fraction of total plasma (<15%) and ascites (<10%) protein, despite a comparable number of EV-like particles per microgram of protein were obtained between the two biofluids **(Figure S1D-E)**. SAX-enrichment depends on electrostatic interactions between the bead-bound ammonium group with the negative macromolecules (e.g. proteins and lipids) comprising the EV membrane^21^. Indeed, negative zeta potential of SAX-enriched EVs from plasma (−15.1±2.1 mV) and ascites (−15.1±2.1 mV) were consistent with previously reported EV ranges of -10mV to -70mV **(Figure 1E)**; moreover, EV-enrichment coincided with depletion of albumin and enrichment of CD9 as the number of washes was increased **(Figure 1F; Figure S1F-G)**. SAX-enriched EVs represent a fraction (∼1%) of the total particles in raw and NC fractions; however, only human EVs were smaller in diameter than particles in raw or NC fractions **(Figure 1G-H, Figure S1C)**. Consistent with previous studies, implementation of the Mag-Net protocol increased proteomic depth >2-fold **(Figure 1I; Table S1)**. In fact, 1020 proteins were uniquely identified in SAX-enriched EV proteomes compared to raw murine ascites, of which >50 of these proteins were associated with vesicle-mediated transport **(Figure 1J)**. Of the 1947 commonly identified proteins, 1179 and 443 differentially expressed proteins (DEP) were enriched or depleted in EV proteomes, respectively **(Figure 1K)**. Secretory autophagosomes and amphisomes are an EV subclass released by cells during periods of microenvironmental stress^36, 37^. Aligned with predicted stressors and the harsh microenvironment of the ascites fluid, murine EVs were distinctly enriched with autophagic proteins ATG3/7 **(Figure 1L)**. Classical EV proteins, including CD9 and TSG101, were enriched >30-fold and coincided with a >8-fold depletion of albumin and tissue factor **(Figure 1L)**. Not surprisingly, several terms related to EV biogenesis or architecture were identified by Jensen Compartment annotations **(Figure S1H)**. GSEA using KEGG_2019 (murine) annotation highlights an enrichment of proteins related to ECM-receptor interactions **(Figure S1I)**, including integrin-binding Thy-1 Cell Surface Antigen (Thy-1/CD90) that has established roles in stem cell and cancer cell biology^38^. Collectively, these results demonstrate the feasibility of SAX EV enrichment, from human and murine ascites.

**Figure 1.**
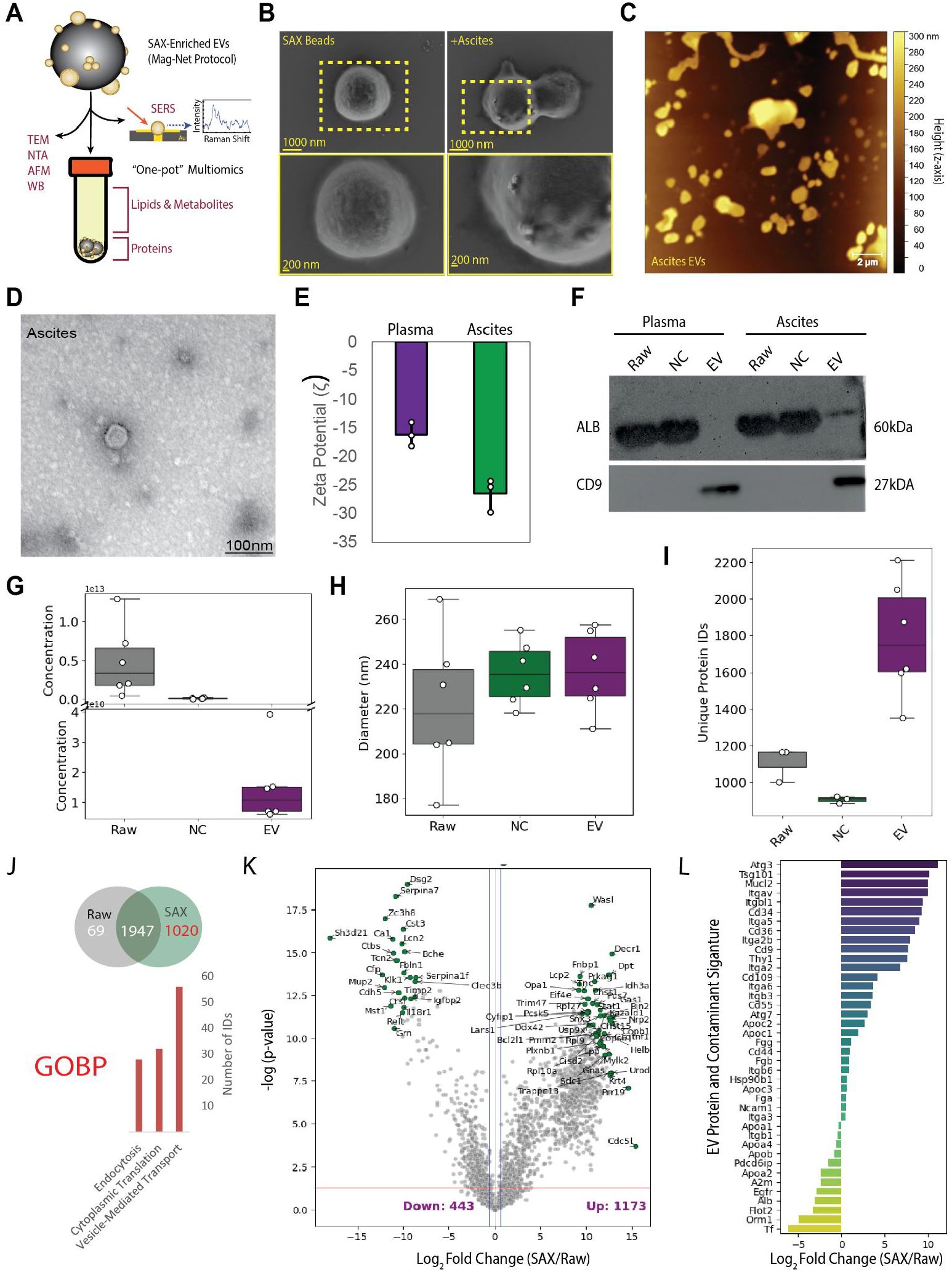
Enrichment and characterization of ascites extracellular vesicles using SAX magnetic beads. EVs were enriched from 100 µL of murine or human ascites fluid using the SAX bead workflow and characterized using orthogonal approaches consistent with MISEV guidelines. **(A)** Schematic overview of SAX bead-based EV enrichment from ascites fluid, enabling downstream EV characterization by TEM, NTA, AFM, and western blotting, as well as single-isolation multiomic profiling, including proteomics, lipidomics, metabolomics, and single-EV chemical fingerprinting by SERS. **(B)** Representative SEM micrographs of SAX beads before and after incubation with ascites fluid, showing EV-like particles associated with the bead surface. **(C)** Representative AFM image of EV-like particles enriched from human ascites and eluted from SAX beads. **(D)** Representative TEM image of human ascites EVs displaying typical vesicular morphology. **(E)** Zeta potential measurements of EVs enriched from human plasma and ascites fluid. **(F)** Western blot analysis of albumin and CD9 in raw input, NC fractions, and SAX-enriched EV fractions from plasma and ascites. SAX enrichment increased CD9 signal while reducing albumin abundance relative to the corresponding raw and NC fractions. **(G, H)** NTA of murine ascites fractions showing particle **(G)** concentration and **(H)** diameter in raw ascites, NC fractions, and SAX-enriched EV fractions. **(I)** Gas-phase fractionation DIA proteomics demonstrated increased proteomic depth in SAX-enriched EV fractions compared with raw ascites and NC fractions. **(J)** Venn analysis showing 1,020 proteins uniquely identified in SAX-enriched EVs relative to raw ascites, with Gene Ontology biological process enrichment highlighting vesicle-mediated transport, cytoplasmic translation, and endocytosis. **(K)** Differential abundance analysis comparing SAX-enriched EVs with raw ascites identified 1,173 proteins enriched and 443 proteins depleted in SAX-enriched EV fractions. **(L)** EV-associated proteins, including Cd9 and Tsg101, were enriched in SAX-enriched murine ascites EVs, whereas abundant soluble or contaminating proteins, including albumin-related proteins, were depleted relative to raw ascites. Data in **(E)** are shown as mean ± SD. Data in **(G–I)** are shown as box plots with median and interquartile range.

### 3.2 Ascites EVs reflect tumor microenvironment and possess a multiomic signature

Within the last decades, EVs have become an attractive source of biomarker for diagnostic and therapeutic management, in part due to their reflection of the tumor proteome and microenvironment^8, 17^. As a proof-of-principle, we compared tumor proteomes to raw ascites and SAX-enriched EVs using GPF-DIA and chemical fingerprinting of single EVs using SERS with fabricated nanohole arrays engineered to samples up to 100 EVs from a drop-casted solution^9^ **(Figure 2A)**. 1627 proteins were commonly identified across the three proteomes; however, 817 proteins were uniquely identified in both EV and tumor proteomes **(Figure 2B; Table S1, S2)**. These 817 proteins were significantly associated with protein translation and RNA metabolism **(Figure S2A)**, which aligns with previous reports of unintentional ribosomal complex enrichment^39^. However, future efforts will need to determine if these signatures represent bonafide EV cargo versus co-enrichment macromolecular complexes forming the EV corona^40^.

**Figure 2.**
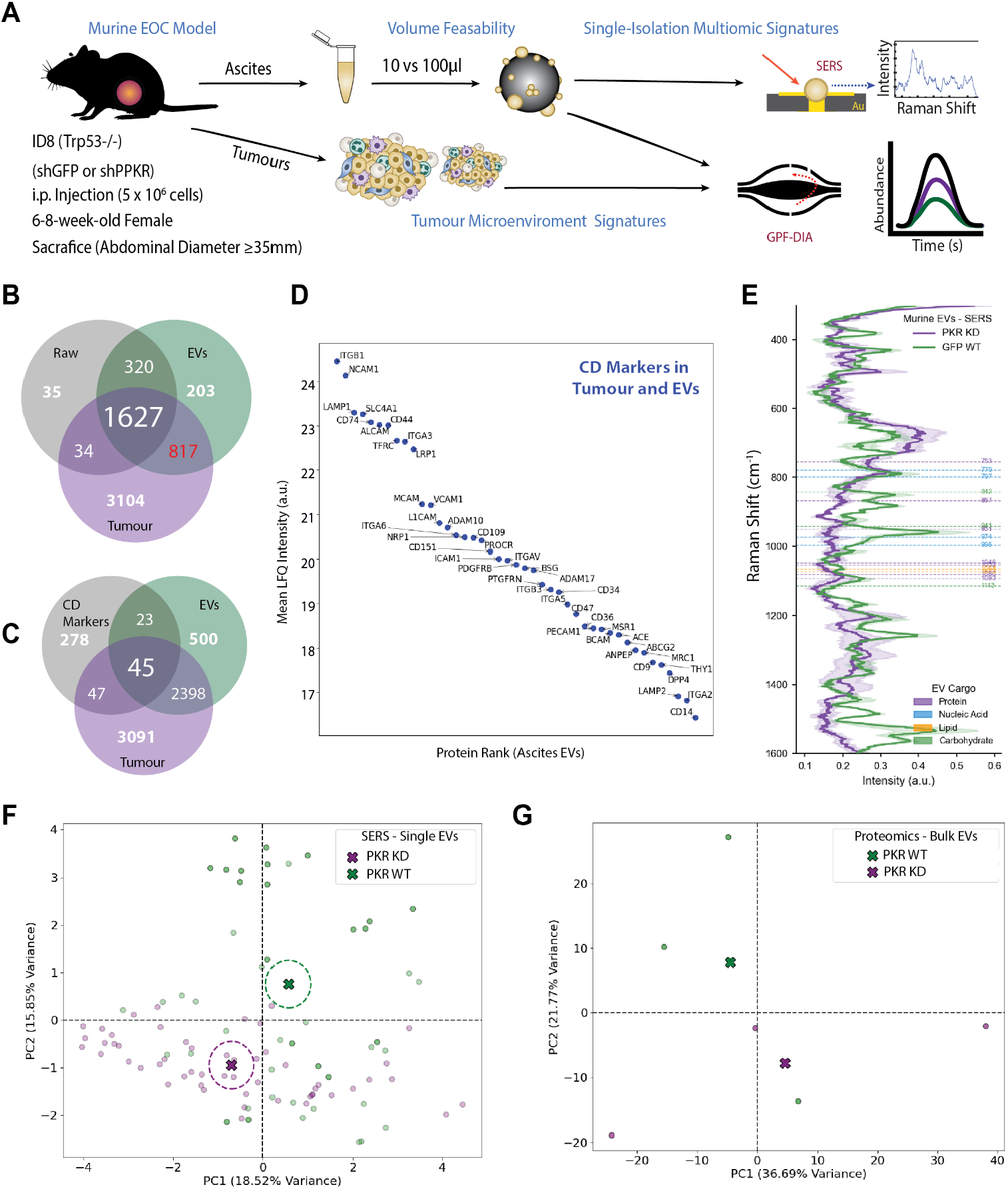
Parallel Profiling of Ascites EVs Using LC-MS/MS and Single-EV SERS. Ascites EVs were profiled using a murine intraperitoneal epithelial ovarian cancer model. ID8 Trp53−/− cells expressing either control shGFP/PKR wild-type (WT) or shRNA-mediated Eif2ak2/PKR knockdown (KD) were injected intraperitoneally, and ascites and tumour tissues were collected at endpoint. **(A)** Workflow schematic illustrating collection of ascites and tumours, SAX-based EV enrichment from ascites fluid, volume feasibility testing using 10 versus 100 µL input, and downstream profiling of SAX-enriched EVs by gas-phase fractionation DIA proteomics and single-EV SERS. Tumour tissues were analyzed in parallel to compare EV proteomes with the tumour microenvironment. **(B)** Venn analysis comparing proteins identified in raw ascites, SAX-enriched EVs, and tumour proteomes. SAX-enriched EVs shared 1,627 proteins with both raw ascites and tumour proteomes, while 817 proteins were detected in both SAX-enriched EVs and tumour tissue but not in raw ascites. **(C)** Venn analysis of cluster of differentiation (CD) markers identified across raw ascites, SAX-enriched EVs, and tumour proteomes. Forty-five CD markers were common to all three fractions, while 23 CD markers were detected in both raw ascites and SAX-enriched EVs but not in tumour proteomes, and 500 CD markers were uniquely detected in SAX-enriched EVs. **(D)** Rank-abundance analysis of CD markers identified in ascites EVs and tumour proteomes, highlighting highly abundant EV-associated markers including ITGB1 and NCAM1. **(E)** Representative SERS spectra from WT and PKR KD murine ascites EVs, showing spectral regions associated with major EV biochemical classes, including proteins, lipids, carbohydrates, nucleic acids, and metabolites. **(F)** Principal component analysis (PCA) of single-EV SERS spectra from SAX-enriched WT and PKR KD ascites EVs. Individual points represent single-EV spectra, and large crossed markers indicate group centroids. **(G)** PCA of bulk EV proteomes generated from SAX-enriched ascites EVs, showing separation between WT and PKR KD EV proteomic profiles.

Given that ascites EVs are enriched in ECM-receptor and focal adhesion components **(Figure 1SH)**, we sought to determine if surface proteins identified on EVs reflect the heterogeneity of parental tumor and infiltrating cell populations within the tumor microenvironment^17^. 45 out 393 CD markers were commonly identified between ascites and tumor, plus an additional 37 and 23 markers identified exclusively in tumors or EVs, respectively **(Figure 2C)**. Amongst commonly identified CD markers, integrin beta 1 (Itgb1/ITGB1) and neural cell adhesion molecule (Ncam/NCAM) were detected at the highest abundance level within ascites EVs **(Figure 2D)**, with both proteins having known roles in OC progression^41, 42^. Furthermore, CD markers identified in both EVs and tumors reflect the cellular heterogeneity present within ascites fluid; including, endothelial and platelet marker (Pecam1), fibroblast marker (Procr), and monocyte/macrophage markers CD14 and Macrophage Mannose Receptor C-Type 1 (Mrc1/MRC1) to highlight a few. CD markers exclusively identified in ascites EVs but not tumor proteomes were associated with markers of platelets, including CD41a/Itga2b, CD42b/Gp1ba, CD42d/Gp5, and CD62L/Sell; fibroblasts, including CD140a/Pdgfra, CD325/Cdh2, CD331/Fgfr1, and CD333/Cd33; leukocytes, including CD85a/Lilrb3 and CD115/Csf1r; and perivascular cells, including CD93/Cd93 and CD248/Cd248. Another important observation was our inability to identify Mucin-1 (Muc1) or Mucin-16 (Muc16) in either murine EVs or tumor proteomes, suggesting these markers may persist below the limit of detection or reflect physiological differences between murine models and primary human samples^43^. In support of the former, epithelial protein Epcam/EPCAM, a surrogate marker for malignant cancer cells, was identified in tumor proteomes but not in the proteomes of EVs **(Table S2)**. These results align with evidence from previous studies that the majority of cells present within the ascites microenvironment are primarily not tumor-derived, but are infiltrating stromal or immune cell populations^44, 45^. Future studies will need to consider dynamic proteome range of ascites and disease-relevance of the selected model^46^ prior to experimental interpretation.

As additional proof-of-concept, we sought to determine if murine ascites EVs carry multiomic signatures, that are reflective of physiological or disease states; including tumor burden. For context, mice receiving shPKR ID8 cells displayed decreased tumor burden compared to shGFP controls **(Figure S2B)** by mechanisms established in ongoing work outside the focus of this study. We have previously demonstrated that a single-EV isolation using SEC from 100µl of blood plasma is sufficient to support both SERS and proteomic analysis of EVs^10^. Accordingly, we show that a single SAX EV-isolation from 100µl or 10µl of starting ascites volume is sufficient to capture a multiomic EV signatures by splitting EV-bound beads 1:1 after gently resuspending beads in solution^28^ **(Figure S2C-E; Table S3)**. Chemical fingerprints obtained with SERS can include spectral peaks associated with EV cargo including proteins, nucleic acids, and metabolic substrates **(Figure 2E)**. For example, the tryptophan ring (753 cm^−1^), proline stretching (1046 cm^−1^) and C-C collagen backbones (867 cm^−1^ and 1080 cm^−1^). Interpretation of SERS spectra and GPF-DIA data from a single isolation, using principal component analyses, exhibited distinguishable signatures at the single-EV level using SERS **(Figure 2F)** and global EV proteome using GPF-DIA **(Figure 2G)**. Importantly, these distinguishable SERS and proteomic signatures were retained with input volume was lowered to 10µl of ascites fluid; however, less single-EV events were obtained **(Figure S2F-G)**. Collectively, we provide evidence of the feasibility of SAX beads to support multiomic EV profiling utilizing parallel SERS and GPF-DIA proteomic profiling.

### 3.3 SAX EV enrichment enhances proteomic depth compared to UC

We next sought investigate the benefits of SAX-enrichment compared established EV isolation methods, such as UC. We focused these comparisons on a single patient a 59-year-old patient with Stage 1b borderline mucinous carcinoma, in technical triplicate to limit biological variability observed across independent donors **(Figure S3A; Table S4)**. Using 100µl as our baseline input volume, EV-like particles were identified in both SAX or UC enrichment strategies using AFM **(Figure 3A)**; however, we also observed a decreased background in SAX versus UC EV isolations. Elevated background during AFM can indicate residual salts, free proteins, or other liable macromolecules that may not be effectively depleted during EV enrichment. EVs enriched by SAX were statistically smaller than UC or raw samples (85.0 ± 2.0 vs 107.3 ± 9.3 vs 129.67 ± 10.26 nm diameter; **Figure 3B**) and at a reduced concentration (5.3E8 ± 5.7E7 vs 1.5E9 ± 3.6E8 vs 2.0E10 ± 1.5E9 particles/µl; **Figure 3C**). Normalization of EV concentration to total protein recovery indicate a similar EV enrichment efficiency **(Figure S3B)**; however, distinct proteomic signatures exist between SAX- and UC-enriched EVs **(Figure 3D; Figure S3C-D)**. We identified 1049 proteins unique to SAX and UC isolations that were determined to have a statistically pronounced EV signature **(Figure 3E, Table S5)**.

**Figure 3.**
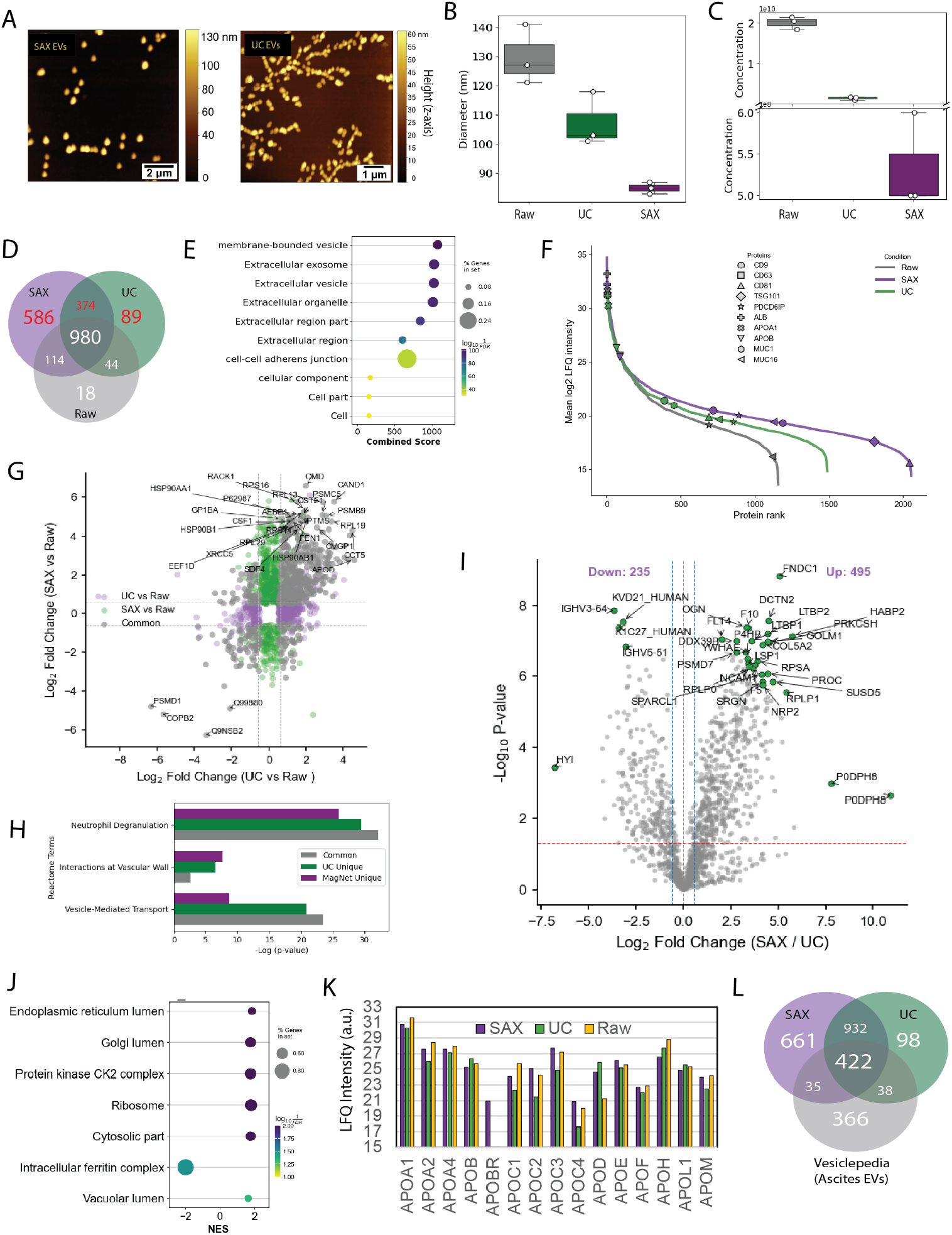
Comparison of SAX versus Ultracentrifugation for the Isolation of EVs from Human Ascites Fluid. Human ascites fluid from a 59-year-old donor with stage IB mucinous ovarian cancer was used to compare SAX magnetic bead enrichment with differential UC (130,000 × g). **(A)** Representative AFM images of EV-like particles enriched from ascites fluid by SAX or UC. **(B**,**C)** NTA comparing particle **(B)** diameter and **(C)** concentration in raw ascites, UC-enriched EVs, and SAX-enriched EVs. **(D)** Venn analysis of proteins identified by gas-phase fractionation DIA proteomics across raw ascites, SAX-enriched EVs, and UC-enriched EVs. SAX enrichment identified 586 unique proteins, while SAX and UC together identified 1,049 proteins not detected in raw ascites. **(E)** Jensen compartment enrichment analysis of proteins detected uniquely in SAX- and/or UC-enriched fractions relative to raw ascites, highlighting significant enrichment of extracellular vesicle-, exosome-, membrane-bound vesicle-, and extracellular organelle-associated annotations. **(F)** Protein rank-abundance plot showing increased proteomic depth and expanded dynamic range in SAX- and UC-enriched EV fractions compared with raw ascites. Canonical EV-associated proteins and common plasma/lipoprotein-associated proteins are indicated. **(G)** Comparison of protein abundance changes in SAX- and UC-enriched EV fractions relative to raw ascites, highlighting shared and method-specific differentially abundant proteins. **(H)** Reactome pathway enrichment analysis of proteins commonly enriched or uniquely enriched by SAX or UC, including neutrophil degranulation, interactions at the vascular wall, and vesicle-mediated transport. **(I)** Differential abundance analysis comparing SAX-enriched and UC-enriched EV proteomes, identifying 495 proteins enriched in SAX-EVs and 235 proteins enriched in UC-EVs. **(J)** Gene set enrichment analysis comparing SAX-EVs with UC-EVs, showing enrichment of luminal and intracellular compartment-associated proteins in SAX-enriched EV fractions. **(K)** Mean LFQ intensity of apolipoproteins and lipoprotein-associated proteins across raw ascites, SAX-enriched EVs, and UC-enriched EVs. **(L)** Overlap of proteins identified in SAX-enriched EVs and UC-enriched EVs with Vesiclepedia proteins annotated from ascites EV datasets. Data in **(B**,**C)** are shown as box plots with median and interquartile range.

Indeed, both isolation methods increased proteomic depth when compared to raw ascites fluid including a considerable number of common DEPs **(Figure 3F; Figure S3E-F)**. Relative to raw ascites proteomes, SAX- and UC-enriched DEPs were enriched with components of vesicle transport, neutrophil degranulation, and interactions at the vascular wall **(Figure 3G-H)**. Furthermore, unique cellular compartments were enriched by each method **(Figure 3SG)**. While both UC-enriched and SAX-enriched EV proteomes were commonly associated with blood microparticle and extracellular matrix components, an enrichment of immunoglobulin complexes were observed with SAX-enriched EVs **(Figure S3G)**. Previously studies have observed that immunoglobin complex can bind form the corona of biofluid EVs^47, 48^. These observations require further investigation since differential analysis uncovered a total of 914 DEPs between isolation methods **(Figure 3I)**. GSEA using GOCC uncovered a significant enrichment of ribosomal subunit and organelle lumens in SAX-enriched EVs **(Figure 3J)**. These results align with previous studies comparing EV isolation methods which have observed distinct proteomes while retaining an ISEV minimal EV signature^35, 49^.

Next, we directly compared each isolation method by focusing on classical EV markers in addition to albumin and lipoprotein contamination. Despite SAX beads providing increased proteomic depth compared to UC **(Figure S3C)**, albumin was increased by >2-fold in SAX-enriched EV proteomes and EV marker CD81 was enriched within UC proteomes **(Figure S3H)**. This was in contrast to EV markers (i.e. CD9 and ALIX), OC-relevant biomarkers (i.e. MUC1 and MUC16), and fibronectin (FN1) found to be enriched or comparable in SAX-EV proteomes **(Figure S3H)**. Fibronectin is a known component of the EV corona and elicits signaling activity during OC metastasis at the peritoneal interface^50-52^. Importantly, both SAX and UC did not enrich for lipoproteins relative to raw ascites fluid, indicating effective removal of bulk lipoproteins by both isolation strategies **(Table S4)**. However, each isolation methods revealed unique lipoprotein signatures, including the exclusive detection of apolipoprotein B Receptor (APOBR) in SAX-enriched EVs and elevated Apolipoprotein D (APOD) in both SAX and UC-enriched EVs relative to Raw **(Figure 3K)**. APOD has been reported as EV cargo mediating intercellular communication in the nervous system^53^ and identified as prospective biomarker of non-small cell lung cancer^54^. On the other hand, APOBR is a plasma membrane embedded protein, however there is an absence of literature regarding its role in ovarian cancer or extracellular vesicle cargo^55^. Several HDL-associated apolipoproteins (including APOA1, APOA2, APOM, and APOC family members) were modestly enriched by SAX compared to UC, indicating that residual lipoprotein contamination following SAX enrichment is predominantly HDL-like rather than LDL/VLDL-like. Notably, we provide >2000 proteins within SAX- or UC-enriched EVs not previously annotated within Vesiclepedia **(Figure 3L, Figure S3H)**. While both methods provide cost-effective means for EV isolations, our results indicate that SAX-enrichment provides superior proteomic depth and improved EV purity compared to UC.

### 3.6 Combinatorial or sequential EV enrichment does not increase proteomic depth

While ignoring the increased logistical burden of implementing two isolation methods in series, we sought to assess the benefits of dual enrichment workflows **(Figure 4A)**. Interestingly, proteomes of EVs isolated by sequential combinations of Mag-Net/SAX and UC were more similar to each other than to either method alone and combinatorial enrichment strategies did not increase proteomic depth **(Figure 4B-C, Figure S4A)**. In fact, we observed the highest number of proteins identified when the Mag-Net protocol was implemented alone **(Figure S4A)**. Only 126 additional proteins were exclusively detected with dual isolations. However, combinatorial enrichment strategies did increase EV purity as determined by levels of EV markers CD9, CD47, and ALIX in junction with decreased albumin **(Figure 4D-E**). Taken together, we can conclude that combinations of UC and Mag-Net workflows do not provide benefits which outweigh logistical requirements.

**Figure 4.**
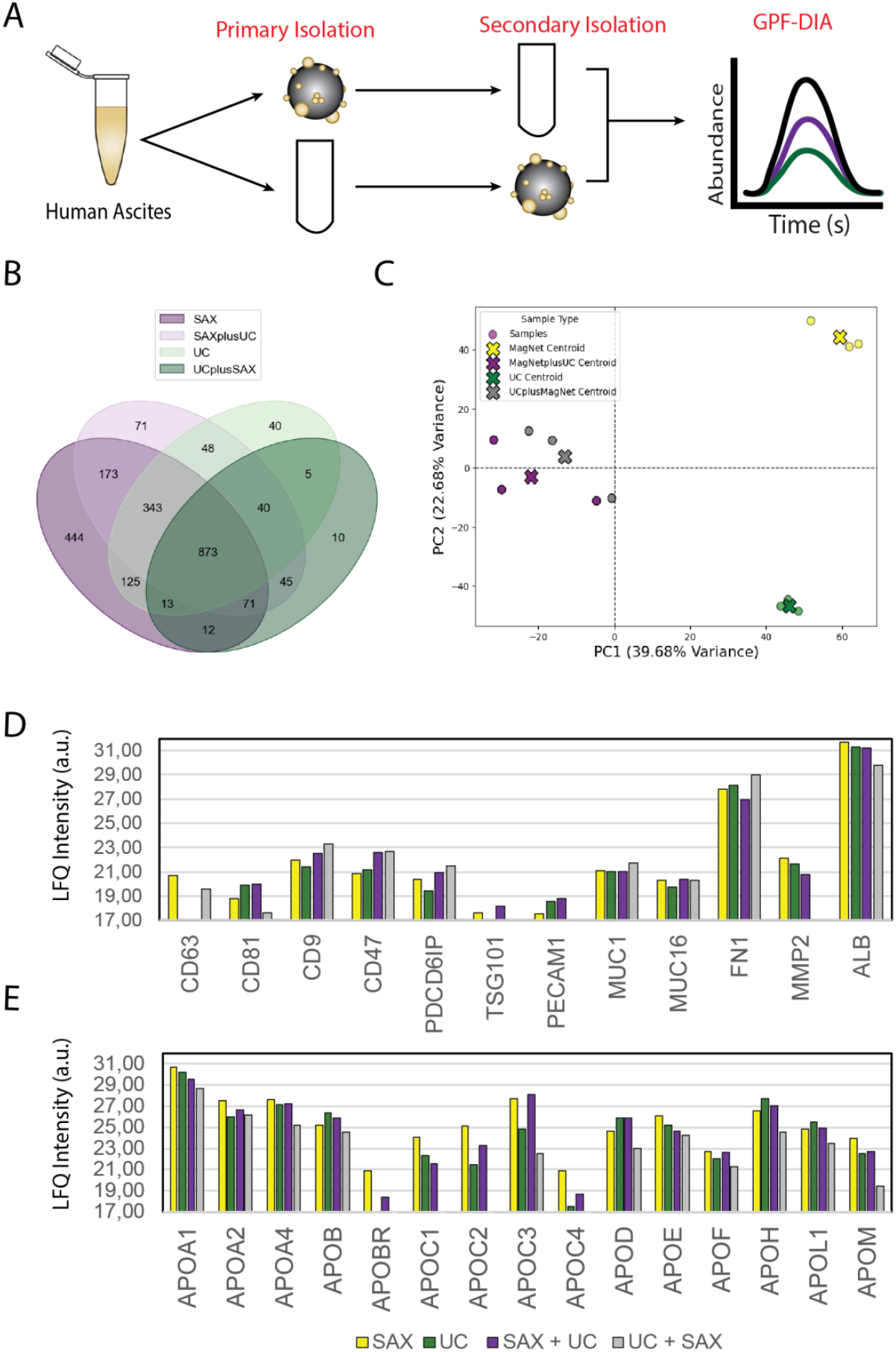
Combinatorial isolation of EVs using SAX and UC does not increase depth of proteomic analysis yet reinforces EV signatures. Human ascites fluid was subjected to single or sequential EV enrichment workflows using SAX magnetic beads, UC, SAX followed by UC, or UC followed by SAX. **(A)** Workflow schematic illustrating primary EV enrichment by SAX or UC followed by secondary enrichment using the reciprocal workflow prior to gas-phase fractionation DIA proteomic analysis. **(B)** Venn analysis comparing proteins identified across single and sequential EV enrichment strategies. A core set of 873 proteins was detected across all workflows, while sequential enrichment did not substantially increase the number of proteins identified relative to single-method enrichment. **(C)** Principal component analysis of EV proteomes generated by single and sequential enrichment workflows. Sequential SAX + UC and UC + SAX proteomes clustered more closely with one another than with either single SAX or UC enrichment, indicating convergence of the EV proteomic profile following combined isolation. **(D)** Mean LFQ intensity values of canonical EV-associated proteins, including tetraspanins, ESCRT-associated proteins, adhesion molecules, and extracellular matrix-associated EV proteins, across SAX, UC, SAX + UC, and UC + SAX workflows. **(E)** Mean LFQ intensity values of apolipoproteins and lipoprotein-associated proteins across the same enrichment workflows.

Understanding the preference of SAX beads for specific EV populations is important to interpreting proteomic results. Thus, we focused on the NC fraction of primary SAX isolations to investigate if SAX beads have preferential enrichment towards a specific EV subpopulation in primary versus secondary isolations. In agreement with our NTA analysis **(Figure 1G-I)**, the complexity of NC is distinctly increased compared to primary EV enrichments. It is likely the NC fraction presents as a mixture of residual EVs, ECM, and undetermined cellular components **(Figure 5A, Figure S4B)**. In order to determine if residual EVs, possessing SAX-compatible biophysical properties, remain in the NC fraction, we performed a secondary isolation using either SAX or UC enrichment protocols followed by GPF-DIA proteomics **(Figure 5B)**. Both SAX and UC enrichment protocols increased proteomic depth compared to the NC fraction alone; moreover, proteomic depths from secondary isolations were comparable the depth of EV proteomes from primary isolations **(Figure 5C)**. This data suggests that EVs from primary and secondary isolations likely carry similar proteomic cargo, and capture is in part a stochastic sample of EVs. In support of this speculation, NC+SAX and NC+UC clustered closer to their respective isolation protocols than each other **(Figure 5D)**, thus these results reinforce the concept that EV proteomes are reflective of isolation methods and opposed to non-specific enrichment^56^. To exemplify this concept, we uniquely identified 758 proteins in SAX and NC+SAX isolations not detected in NC fractions **(Figure 5E)**. Jensen compartments analysis indicated a significant enrichment of EV-related proteins out of the 758 proteins, thus confirming both primary and secondary isolations effectively enrich for EVs **(Figure 5F)**. Furthermore, 1158 and 871 DEPs relative to NC were identified in SAX and NC+SAX EV proteomes **(Figure S4C)**. Cross-analysis of these DEPs identified 944 that were common between primary and secondary SAX isolations **(Figure 5G)**, and these proteins were significantly associated with side of membrane, blood microparticle, and ECM components **(Figure 5H)**. These proteins were also enriched with proteomic components of cytoplasmic translation, blood vessel development, and wound healing **(Figure 5I)**. Secondary isolations were able to increase detectable levels of EV-related proteins relative to NC; however, several HDL-associated lipoproteins were enriched in primary and secondary SAX captures relative to NC, indicating a bias towards HDL-particle contamination without affecting albumin levels **(Figure S4D-F**; **Table S5)**. This observation confirms the capture of residual EVs not isolated by primary SAX-enrichment. Given the similarity of primary and secondary SAX isolations, these results indicate a targeted selection of EVs using SAX beads opposed to stochastic enrichment. Albeit, these data also indicate a saturation of EV binding sites on SAX beads may occur during primary isolations or selectivity based on net surface charge may have occurred. Experimentation outside the scope of this study would be required to elucidate these possibilities.

**Figure 5.**
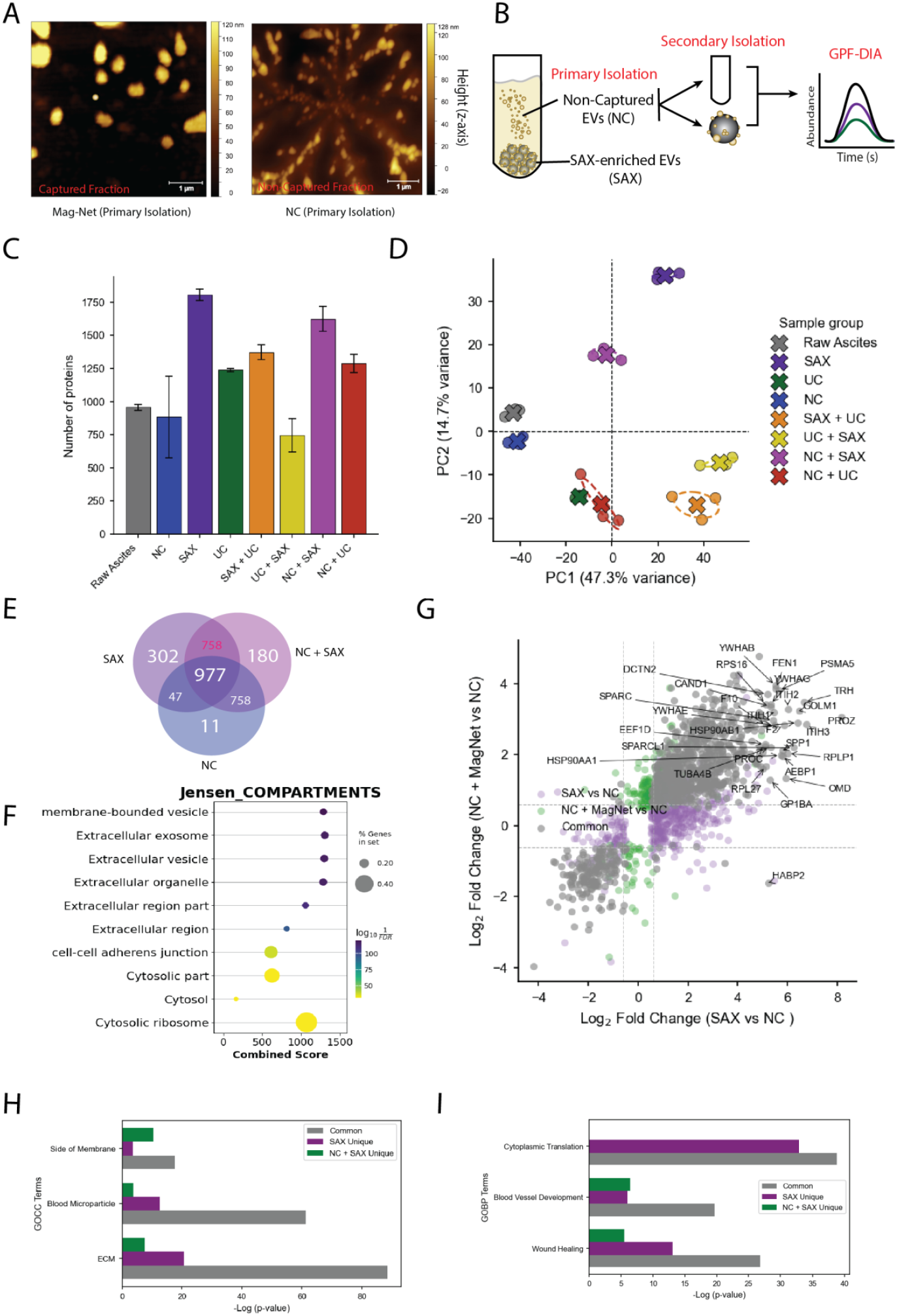
Proteomic Profiling of EVs Enriched from NC fractions of Primary Isolations. To determine whether EV remained in the NC fraction after primary Mag-Net/SAX enrichment, NC fractions were subjected to a secondary isolation step using either SAX magnetic beads or UC, followed by GPF-DIA proteomic analysis. **(A)** Representative AFM images of the primary SAX-captured fraction (left) and the corresponding NC fraction (right). EV-like particles were observed in both fractions. **(B)** Schematic overview of the workflow, in which the NC fraction remaining after primary SAX enrichment was subjected to secondary EV isolation by either UC or SAX prior to proteomic analysis. **(C)** Number of proteins identified across raw ascites, NC, primary SAX, primary UC, sequential SAX + UC, sequential UC + SAX, secondary NC + SAX, and secondary NC + UC fractions. Secondary isolation of NC fractions increased proteomic depth, with NC + SAX yielding more identified proteins than NC + UC. **(D)** Principal component analysisnof proteomic profiles across all isolation strategies. NC + SAX and NC + UC clustered more closely with primary SAX and UC EV proteomes, respectively, than with the NC fraction, indicating recovery of EV-associated proteomic signatures following secondary isolation. **(E)** Venn analysis comparing proteins identified in primary SAX, NC, and secondary NC + SAX fractions. A large number of proteins were shared between primary SAX and secondary NC + SAX fractions, whereas relatively few proteins were unique to NC alone. **(F)** Jensen compartment enrichment analysis of proteins shared between primary SAX and secondary NC + SAX fractions but absent from NC alone, showing significant enrichment for extracellular vesicle- and membrane-bound vesicle-associated annotations. **(G)** Cross-comparison of differential abundance analyses for SAX versus NC and NC + SAX versus NC. This analysis identified 944 commonly enriched proteins, 506 proteins uniquely enriched in primary SAX fractions, and 174 proteins uniquely enriched in secondary NC + SAX fractions. **(H**,**I)** Functional enrichment analysis of common and workflow-specific proteins using **(H)** Gene Ontology Cellular Component (GOCC) and **(I)** Gene Ontology Biological Process (GOBP) annotations.

### 3.5 Enrichment of EVs from minimal ascites fluid

At this point we have established the capacity of SAX beads to effectively capture EVs from human ascites fluid from standard volumes (100µl). Accordingly, we sought to characterize the capture of EVs using a 10x reduction (10µl) in input volume **(Figure S5A)**. As anticipated, total protein recovery and particles per microgram of protein were significantly impacted by reduced input volume from the same donor **(Figure 6A-B)**. On the other hand, EVs isolated from 10µl input volume were larger than EVs isolated from 100µl without an observable effect on zeta potential (100µl; -26.5 ± 2.9 versus 10µl; - 23.40 ± 1.90 mV) between the two input volumes **(Figure 6C-D)**. Similar to murine ascites, we demonstrate the feasibility of multiomic analysis using single-EV SERS and GPF-DIA by splitting beads for parallel analyses **(Figure S6A, Figure 6E-K)**. Indeed, the total number of recorded EV events was reduced when EVs were isolated from 10µl versus 100µl input volume without impacting the spectra intensity for a single-EV recording (**Figure 6E)**. Despite a shared donor, principal component analysis dimensionality reduction displayed a separation of single EVs by input volume across principal component 2 **(Figure 6F)**.

**Figure 6.**
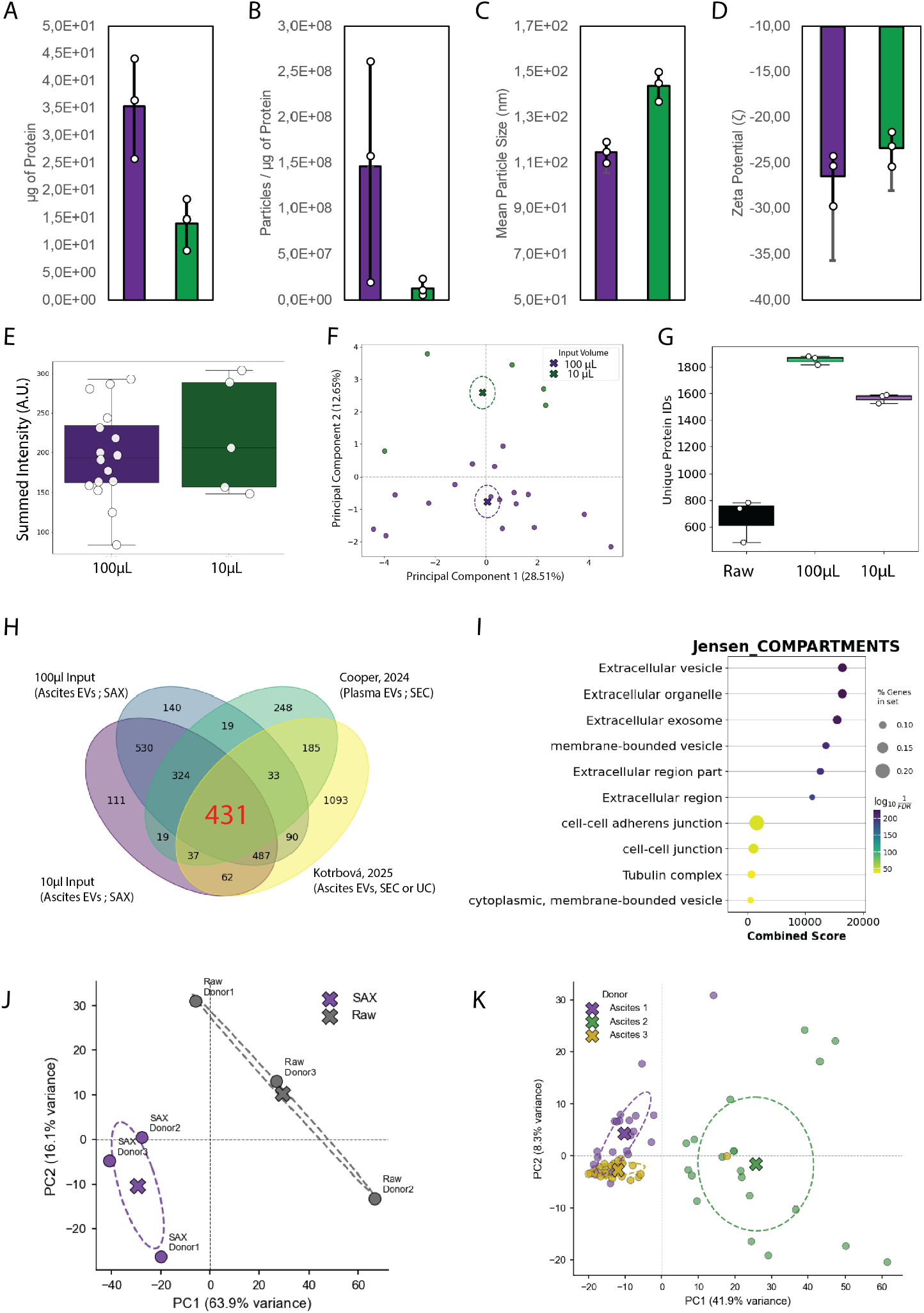
Parallel SERS and Proteomic Profiling of Human Ascites EVs Isolated from Minimal Input Volume. EVs were enriched from human ascites fluid using SAX magnetic beads from either 100 µL or 10 µL of starting material to assess the feasibility of parallel multiomic profiling from minimal sample input. **(A–D)** Comparison of EV yield and biophysical characteristics between input volumes, including **(A)** protein recovery, **(B)** particle-to-protein ratio, **(C)** mean particle size, and **(D)** zeta potential. **(E)** As a proof-of-principle for parallel profiling from a single isolation, single-EV SERS was performed on SAX-enriched EVs obtained from 100 µL or 10 µL input. Summed Raman signal intensity showed a comparable distribution between input volumes. **(F)** Principal component analysis of single-EV SERS spectra derived from 100 µL and 10 µL input volumes. **(G)** Gas-phase fractionation DIA proteomics of SAX-enriched EVs showed greater proteomic depth than raw ascites for both input volumes, with only a modest reduction in the number of proteins identified from 10 µL relative to 100 µL input. **(H)** Venn analysis comparing EV proteomes identified in this study from 100 µL and 10 µL ascites input with previously reported human plasma EV proteomes enriched by SEC and human ascites EV proteomes enriched by SEC or UC. A core set of 431 proteins was shared across all datasets. **(I)** Jensen compartment enrichment analysis of the 431 shared proteins demonstrated significant enrichment for extracellular vesicle-associated annotations. **(J**,**K)** Validation of the bead-splitting workflow using 100 µL ascites from three independent donors with early-stage (FIGO I) high-grade serous carcinoma. **(J)** PCA of proteomic profiles from paired raw ascites and SAX-enriched EV fractions demonstrates donor-specific proteomic signatures and separation of raw versus EV-enriched samples. **(K)** PCA of single-EV SERS spectra from the same donors shows donor-specific clustering of spectral profiles. Data in **(A–D)** are presented as mean ± SD. Data in **(E)** and **(G)** are shown as box plots with median and interquartile range.

Proteomic depth was slightly decreased in EVs from 10µl input volume **(Figure 6G)** where >300 were uniquely identified in one input volume and >500 DEPs were quantified despite sharing a parental donor **(Figure 6G-H; Figure 5SB)**. EV markers CD9 was enriched in 100ul input relative to 10ul input volume, however other EV markers were comparable **(Figure S5C)**. The significance of these DEPs is unclear, however, the influence of input volume on proteomic identification and quantification will need to be taken into consideration during future studies. Comparing proteomic identifications in this study, with our previous study of plasma EVs from early-stage HGSC donors^8^ and a recent study with ascites EVs isolated by UC or SEC from donors with HGSC^17^, we uncovered 431 common proteins enriched with EV-associated proteins according to Jensen Compartments **(Figure 6H-I)**. In this comparison, 791 and 1096 proteins were exclusively identified in SAX- or UC/SEC-enriched EV proteomes, respectively. Finally, using three independent donors with early-stage high-grade serous carcinoma (FIGO I), we validate the feasibility of bead-splitting from 100ul input by identifying OC- and EV-related proteins and retaining measurable single-EV SERS and global proteomic profiles **(Figure 6J-K; Table S6)**. The proteomes of independent donor EVs are distinct **(Figure 6J, Figure S5D)**; yet, consistently provide elevated proteomic depth and were consistently enriched with CD9 and depleted from ALB contamination relative to raw ascites **(Figure S3A; Figure S5E-F)**. Consistent with our previous isolations, we observed an enrichment of translational and luminal proteins with SAX-isolated EVs relative to raw ascites proteomes **(Figure S5G)**. Nonetheless, the number of single EV SERS profiles captured was similar across donors **(Figure S5H)** and Raman signatures of macromolecules were consistent with EV cargo, such as proteins and nucleic acids **(Figure S5I)**. Our future studies are currently implementing these workflows on a cohort of patients suitable for machine learning model training and prospective biomarker discovery efforts.

Current state-of-the-art proteomic facilities include robotic liquid and sample handlers to increase throughput and reproducibility while minimizing the working volume of reactions. In this study, we assess the limitation of manual isolation by testing input volumes ranging between 100µl to 1µl **(Figure 7A)**. Notably, we were able to increase proteomic depth over raw ascites proteomes using a minimum of 2µl or more input volume before proteomic depth was impacted **(Figure 7B-C, Table S6)**. EV markers, such CD9, MUC1, and FN1 and were detected down to 2µl of input volume; however, other markers such as CD81, ALIX, and CD63 required at least 5µl of input volume **(Figure 7C-D)**. Below 100ul input volume, we observed unsupervised clustering around replicated rather than input volume, highlighting the potential of technically variability at low input volumes within the same donor **(Figure S6A)**.

**Figure 7.**
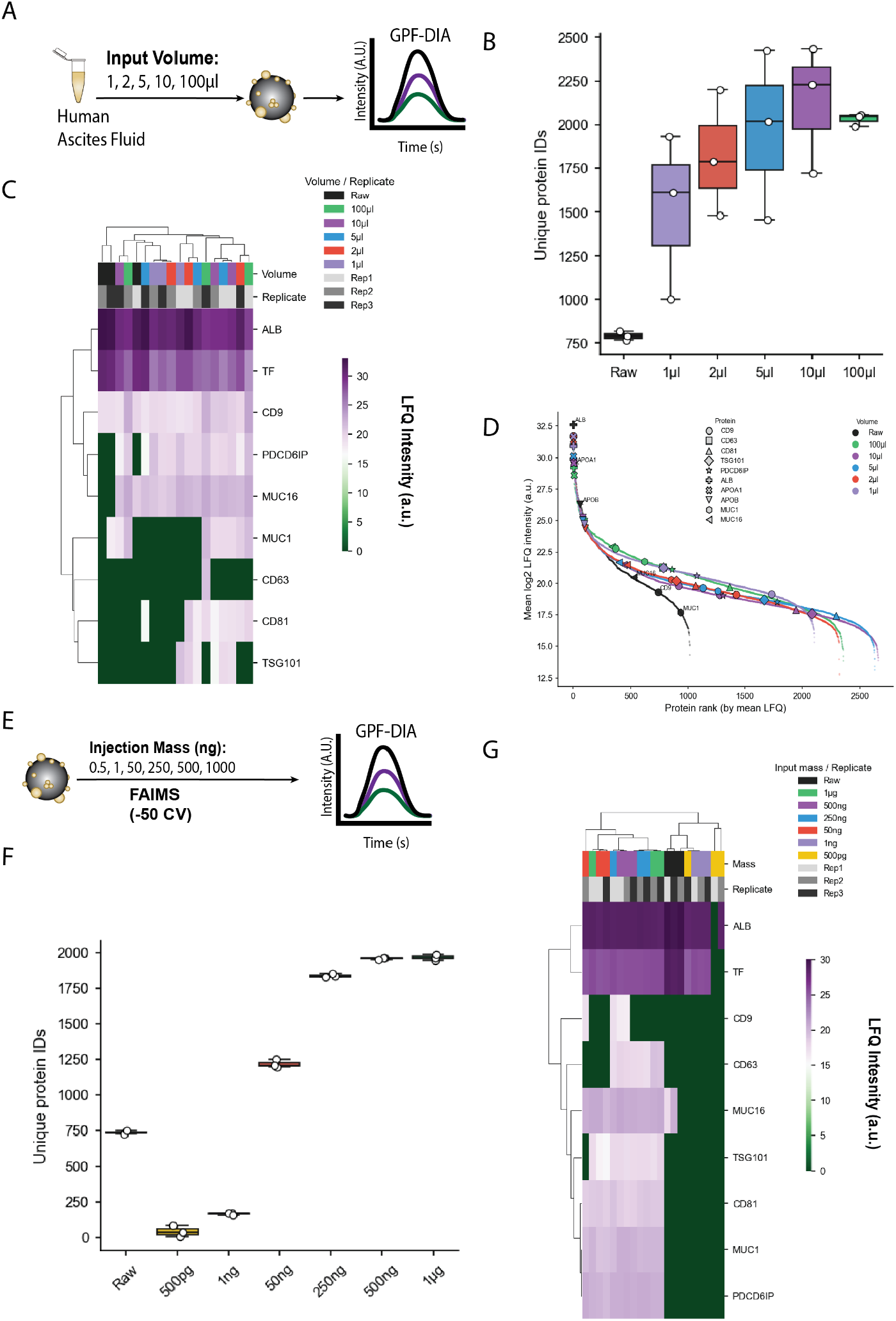
Investigating the minimal input volume or injection mass for robust proteomic profiling of ascites EVs. To establish the lower limits for EV proteomic analysis from human ascites fluid, we evaluated both the minimum ascites input volume required for EV enrichment and the minimum peptide injection mass required for LC– MS/MS analysis. **(A)** Workflow schematic for the input-volume experiment. EVs were enriched from 1, 2, 5, 10, or 100 µL of human ascites fluid using SAX magnetic beads, while maintaining a constant bead-to-protein ratio (1:4). Following digestion, peptides were resuspended in equal volumes of 0.1% formic acid/0.01% DDM and analyzed by GPF-DIA. **(B)** Number of unique protein identifications obtained from raw ascites and from SAX-enriched EVs generated from decreasing input volumes. Although proteome depth declined with reduced starting volume, all EV-enriched fractions yielded more protein identifications than raw ascites. **(C)** Unsupervised clustering and heatmap of LFQ intensities for representative contaminant- and EV-associated proteins across input volumes and replicates. **(D)** Rank-abundance analysis showing mean log2 LFQ intensity as a function of protein rank across raw ascites and SAX-enriched EV samples from each input volume. Lower input volumes showed increased relative contribution of abundant contaminants and reduced EV-marker intensity; however, all EV-enriched fractions exhibited greater proteomic depth and dynamic range than raw ascites. **(E)** Workflow schematic for the injection-mass experiment. Using EVs enriched from 10 µL ascites input, peptide injections ranging from 500 pg to 1 µg were analyzed by GPF-DIA with FAIMS (CV = −50) to determine the minimum injection mass required for sensitive EV proteomics. **(F)** Number of unique protein identifications obtained across peptide injection masses. Relative to raw ascites digests, EV-enriched samples yielded increased protein identifications with injection masses as low as 50 ng. **(G)** Unsupervised clustering and heatmap of LFQ intensities for representative EV- and contaminant-associated proteins across injection masses and replicates. Data in (B) and (F) are shown as box plots with median and interquartile range.

The inclusion of FAIMS analysis can increase the sensitivity of LC-MS/MS analysis and provide confident spectrum-peptide identification due to removal of uninformative ions by optimizing the CV(s) implemented. We selected a CV of –50, based on our previous studies^10^ and removal of air-born contaminants in our facility before injection of decreasing input mass from a single input volume of 10µl **(Figure 7E)**. Compared to 1µg of raw ascites at CV=-50, we show that 50ng is sufficient to increase proteomic depth **(Figure 7F; Table S7);** however, we were only able to identify EV markers (e.g. ALIX, CD9, CD81, and CD63) when at least 250ng was injected **(Figure 7G, Figure S6B)**. Unlike input volume, we observed consistent unsupervised clustering based on injection mass, rather than technical variability between injections **(Figure S6C)**. Based on our previous NTA analysis, we estimate that 50ng of injection material represents 1/20th of a microlitre input volume. This can be extrapolated to roughly 3.0×10^5^ EVs or 30% the volume of a 10µm cell. Collectively, this study rates the utility of SAX beads for low-input ascites volumes and sets the foundation for future studies to identify novel biomarkers across ascites-producing cancers.

## 4 Conclusions

Looking forward, efforts to expand the “multiomic” toolbelt will include lipidomic and transcriptomic analyses that may offer a more comprehensive understanding of EV origin and functions in tumorigenesis^12, 57, 58^. Refining the Mag-Net protocol to accommodate low input volumes and integrating robotic automation could enhance its clinical applicability^59^, particularly in low-yield or minimally invasive samples. The original Mag-Net protocol was optimized for use on a KingFisher^24^, in comparison to manual pipetting throughout this study while handling SAX beads. We would anticipate improved depth and reproducibility using robotic systems; however, our study still exemplifies the lower end of volume for most commercial liquid handlers (i.e. 1µl). We show the feasibility of bead-splitting to support multiomic profiling, including single-EV analyses that can support the development machine learning models^10^. This study also highlights limitations of the SAX-enrichment, such as considerable HDL-protein contamination, enrichment of ribosomal complexes, and a limited ability to further purify EV-specific subpopulations from primary isolations (e.g. UC). Ultimately, we foresee SAX-based EV enrichment strategies isolation as a robust tool for multiomic profiling; and, to advance our ability to monitor disease progression, therapeutic response, and recurrence risk in patients with OC.

## Author contributions

**T.T.C-** Conceptualization, Data Curation, Formal Analysis, Investigation, Methodology, Visualization, Writing-Original Draft, Writing-Review-Editing; **L.V.-** Conceptualization, Methodology, Formal Analysis, Investigation, Writing-Original Draft; **F.A.-** Methodology, Investigation, Writing-Original Draft; **O.F.J.H, C.D**., **R.M**., **T.P.A.J, C.W**., **& T.R**.- Methodology, Investigation; **D.B**., **S.A.A, T.G.S, A.G.-** Supervision, Resources, **F.L-L.-** Supervision, Resources, Writing-Review-Editing; **G.A.L & L.M.P.-** Supervision, Resources, Writing-Review-Editing, Project Administration

## Data availability

LC-MS/MS files were uploaded to PRIDE (PXD068766). Non-proteomic data is available upon request.

## Acknowledgements

We would like to thank members of the Canadian Society for Extracellular Vesicles (CanSEV) for their feedback their inaugural symposium. We would also like to thank members of the Cooper laboratory, Sophie-Anne Blanchet, Ph.D. and Alexane Valence, for there technical assistance, research coordination, and critique during the final revision of manuscript.

