## Supplementary material for "Isolation of Extracellular Vesicles from Minimal Volume Ascites Fluid Using Strong Anion Exchange Beads": Figure S1-S6

Supplemental Figures

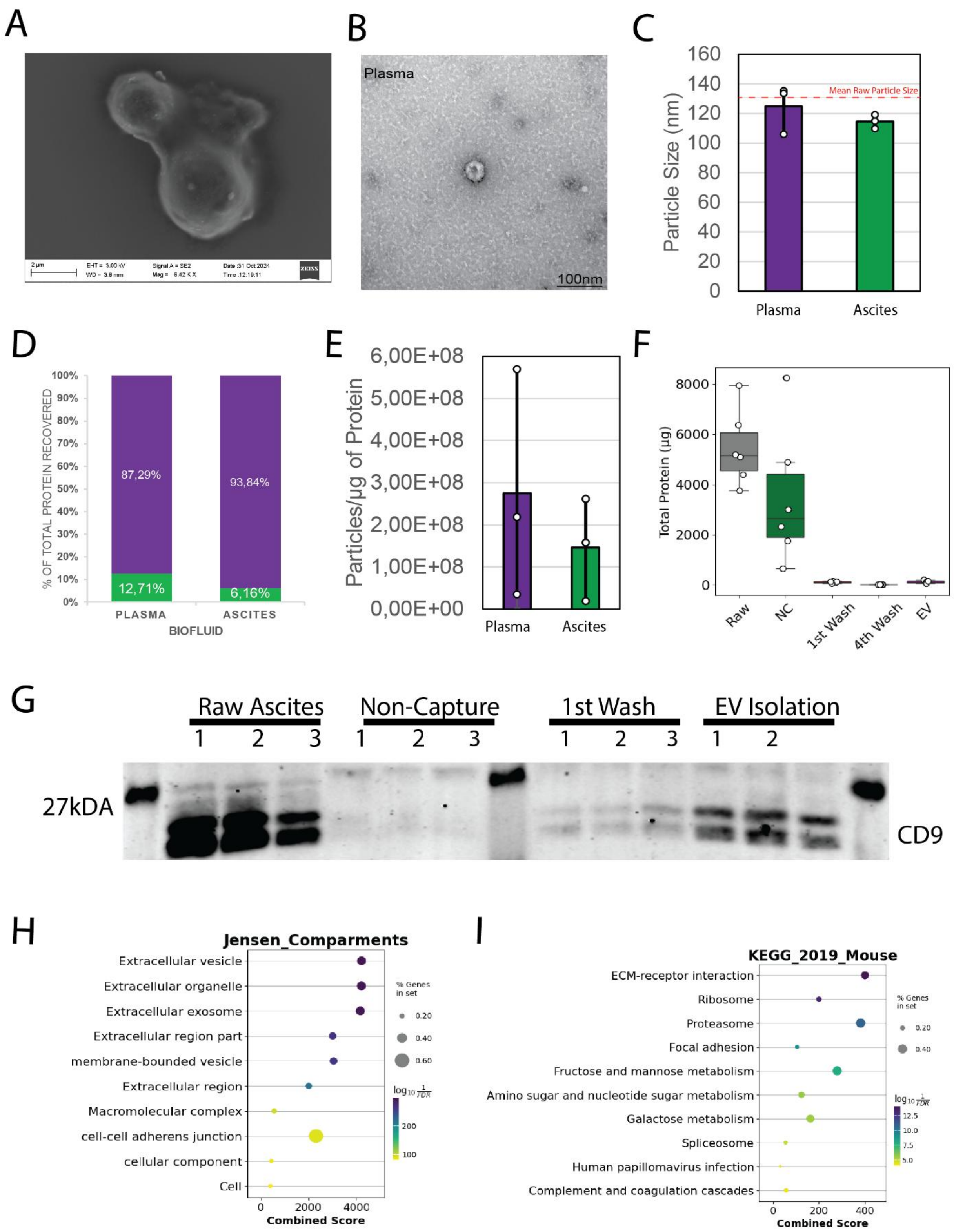

Supplemental Figures

**Figure S1. SAX is effective at enriching EVs from murine and human ascites fluid.** *SAX magnetic beads were evaluated EV enrichment from plasma and ascites fluid using orthogonal EV characterization approaches.* ***(A)*** *Representative SEM image showing bead-associated material and web-like microstructures retained between SAX beads after extensive washing.* ***(B)*** *Representative TEM image of SAX-enriched plasma EVs displaying vesicular morphology.* ***(C)*** *NTA comparing the particle size of SAX-enriched EVs from plasma and ascites fluid. The red dashed line indicates the mean particle size in the corresponding raw ascites.* ***(D)*** *Percentage of total protein recovered in SAX-enriched EV fractions relative to the corresponding raw plasma or ascites input.* ***(E)*** *Particle-to-protein ratio of SAX-enriched EVs isolated from plasma and ascites fluid.* ***(F)*** *Total protein recovered across the SAX enrichment workflow, including raw murine ascites, non-captured fraction, first wash, fourth wash, and final EV-enriched fraction.* ***(G)*** *Western blot analysis of CD9 across raw ascites, non-captured fractions, wash fractions, and final SAX-enriched EV isolates, demonstrating retention and enrichment of CD9-positive EVs following serial washing.* ***(H,I)*** *Enrichment analysis of proteins increased in SAX-enriched EVs relative to raw ascites proteomes using* ***(H)*** *Jensen compartment annotations and* ***(I)*** *KEGG 2019 mouse pathway annotations, highlighting extracellular vesicle/exosome-associated compartments and EV-relevant biological pathways. Data in* ***(C,E)*** *are presented as mean ± SD. Data in* ***(F)*** *are shown as box plots with median and interquartile range.*

Supplemental Figures

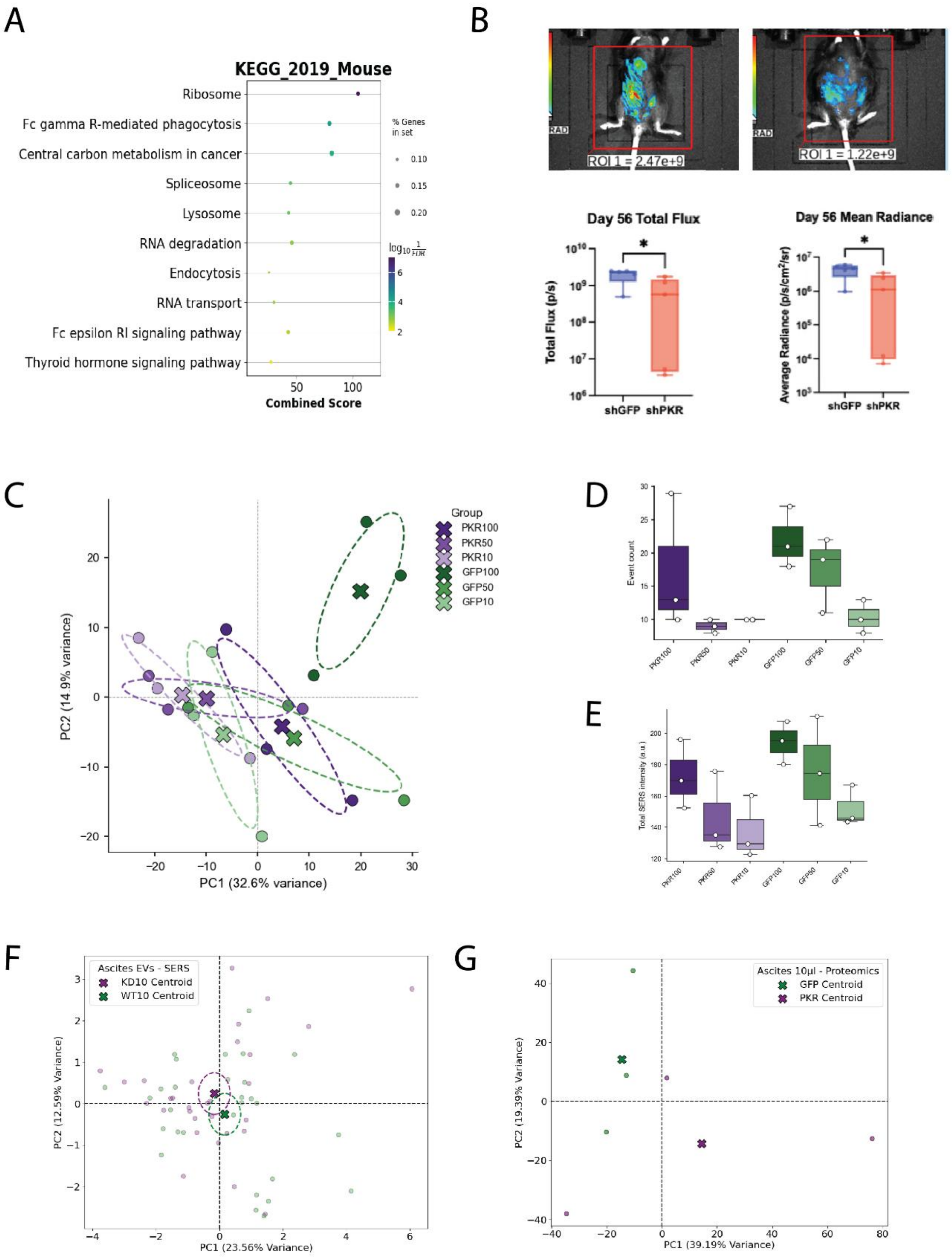

**Figure S2. Ascites EVs are reflective of tumour microenvironment and disease state.** *Ascites EVs enriched by SAX magnetic beads were evaluated for tumour-associated proteomic and single-EV SERS signatures in a murine intraperitoneal epithelial ovarian cancer model.* ***(A)*** *KEGG 2019 mouse pathway enrichment analysis of proteins enriched in SAX-EVs relative to tumour proteomes, highlighting pathways associated with ribosomes, RNA processing, endocytosis, lysosomes, and immune-related signalling.* ***(B)*** *Representative bioluminescence images and quantification of tumour burden at day 56 in mice bearing shGFP control or shPKR tumours, shown as total flux and mean radiance.* ***(C)*** *Principal component analysis of single-EV SERS spectra obtained from SAX-enriched ascites EVs isolated from different starting input volumes: 100, 50, and 10 µL.* ***(D,E)*** *Quantification of single-EV SERS acquisition metrics across input volumes, including* ***(D)*** *total EV event counts and (E) summed SERS intensity.* ***(F)*** *PCA of single-EV SERS spectra from SAX-enriched ascites EVs comparing control and PKR knockdown groups using 10 µL ascites input.* ***(G)*** *PCA of GPF-DIA proteomes generated from SAX-enriched ascites EVs isolated from 10 µL ascites input, comparing control and PKR knockdown groups. Data in* ***(D,E)*** *are shown as box plots with median and interquartile range.*

Supplemental Figures

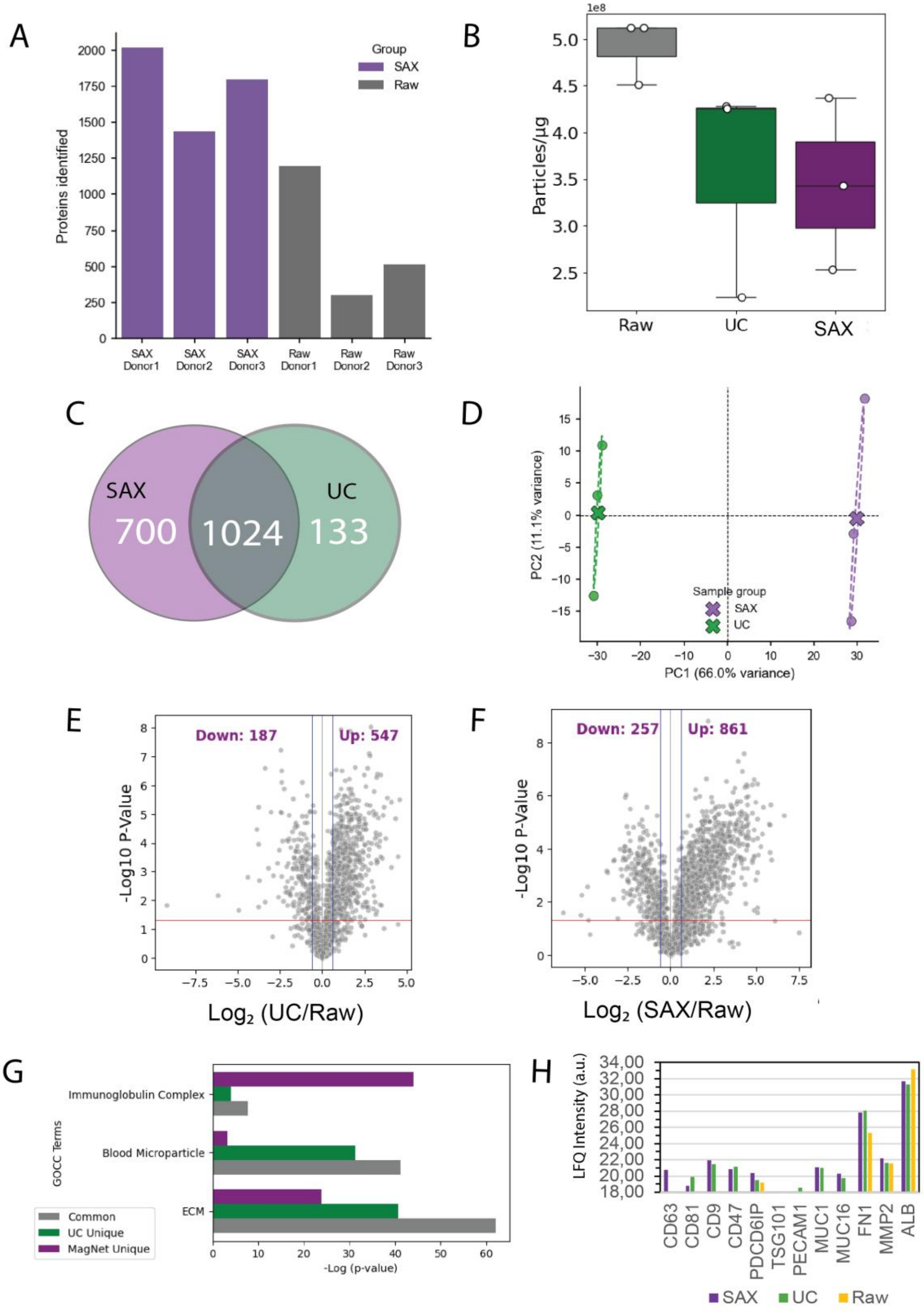

**Figure S3. SAX provides increased proteomic depth relative to UC.** *SAX magnetic bead enrichment and UC were compared for EV proteomic profiling from human ascites fluid.* ***(A)*** *Number of proteins identified in paired raw ascites and SAX-enriched EV fractions from three independent donors, showing increased proteomic depth following SAX enrichment.* ***(B)*** *Particle-to-protein ratio of raw ascites, UC-enriched EVs, and SAX-enriched EVs.* ***(C)*** *Venn analysis comparing proteins identified in SAX- and UC-enriched EV proteomes, showing 1,024 proteins common to both workflows, 700 proteins uniquely detected following SAX enrichment, and 133 proteins uniquely detected following UC enrichment.* ***(D)*** *Principal component analysis of SAX- and UC-enriched EV proteomes, showing separation between isolation workflows along PC1.* ***(E,F)*** *Differential abundance analysis comparing* ***(E)*** *UC-enriched EVs versus raw ascites and* ***(F)*** *SAX-enriched EVs versus raw ascites. UC enrichment identified 547 significantly increased and 187 decreased proteins relative to raw ascites, whereas SAX enrichment identified 861 significantly increased and 257 decreased proteins****.*** ***(G)*** *Gene Ontolo gy Cellular Component enrichment analysis of proteins common to both workflows or uniquely detected by UC or SAX enrichment, highlighting extracellular matrix, blood microparticle, and immunoglobulin complex-associated annotations.* ***(H)*** *EV signatures represented by non-imputed LFQ intensities.*

Supplemental Figures

d

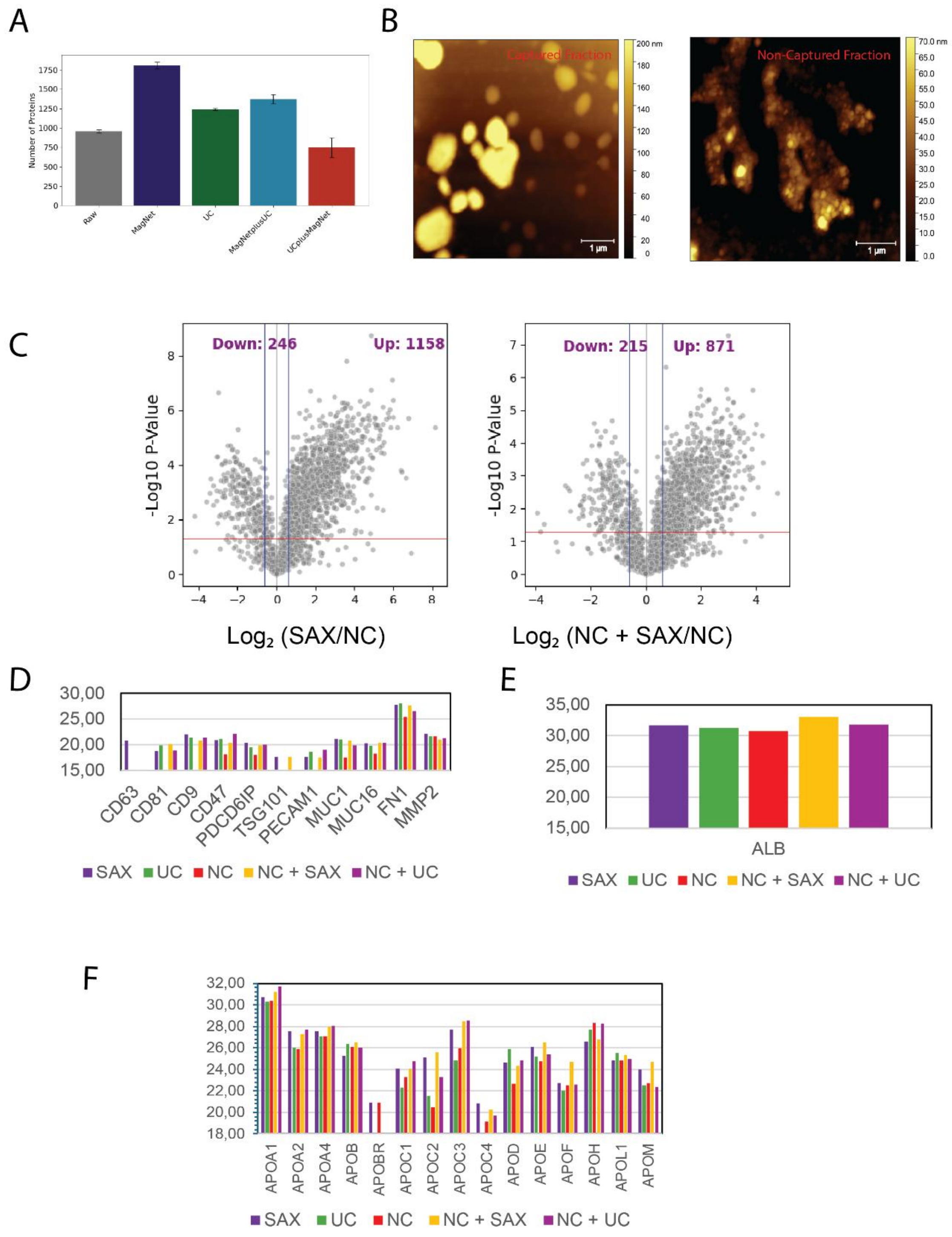

**Figure S4. SAX and UC enrich EVs not captured by primary SAX isolations.** *Secondary enrichment strategies were evaluated to determine whether EV-associated material remained in the NC fraction following primary SAX/Mag-Net isolation.* ***(A)*** *Number of proteins identified across raw ascites, primary Mag-Net/SAX-enriched EVs, primary UC-enriched EVs, Mag-Net followed by UC, and UC followed by Mag-Net workflows.* ***(B)*** *Representative AFM images of the primary Mag-Net/SAX-captured fraction and corresponding NC fraction, showing EV-like particles in both fractions. Albeit, the NC appears much more heterogenous with the presence of non-EV structures.* ***(C)*** *Differential abundance analysis comparing primary SAX-enriched EVs versus NC fractions and secondary NC + SAX fractions versus NC fractions. Primary SAX enrichment identified 1,158 increased and 246 decreased proteins relative to NC, whereas secondary NC + SAX enrichment identified 871 increased and 215 decreased proteins relative to NC.* ***(D)*** *Mean LFQ intensity values of EV-associated proteins across SAX, UC, NC, NC + SAX, and NC + UC fractions.* ***(E)*** *Mean LFQ intensity of albumin across the same fractions.* ***(F)*** *Mean LFQ intensity values of apolipoproteins and lipoprotein-associated proteins across SAX, UC, NC, NC + SAX, and NC + UC fractions.*

Supplemental Figures

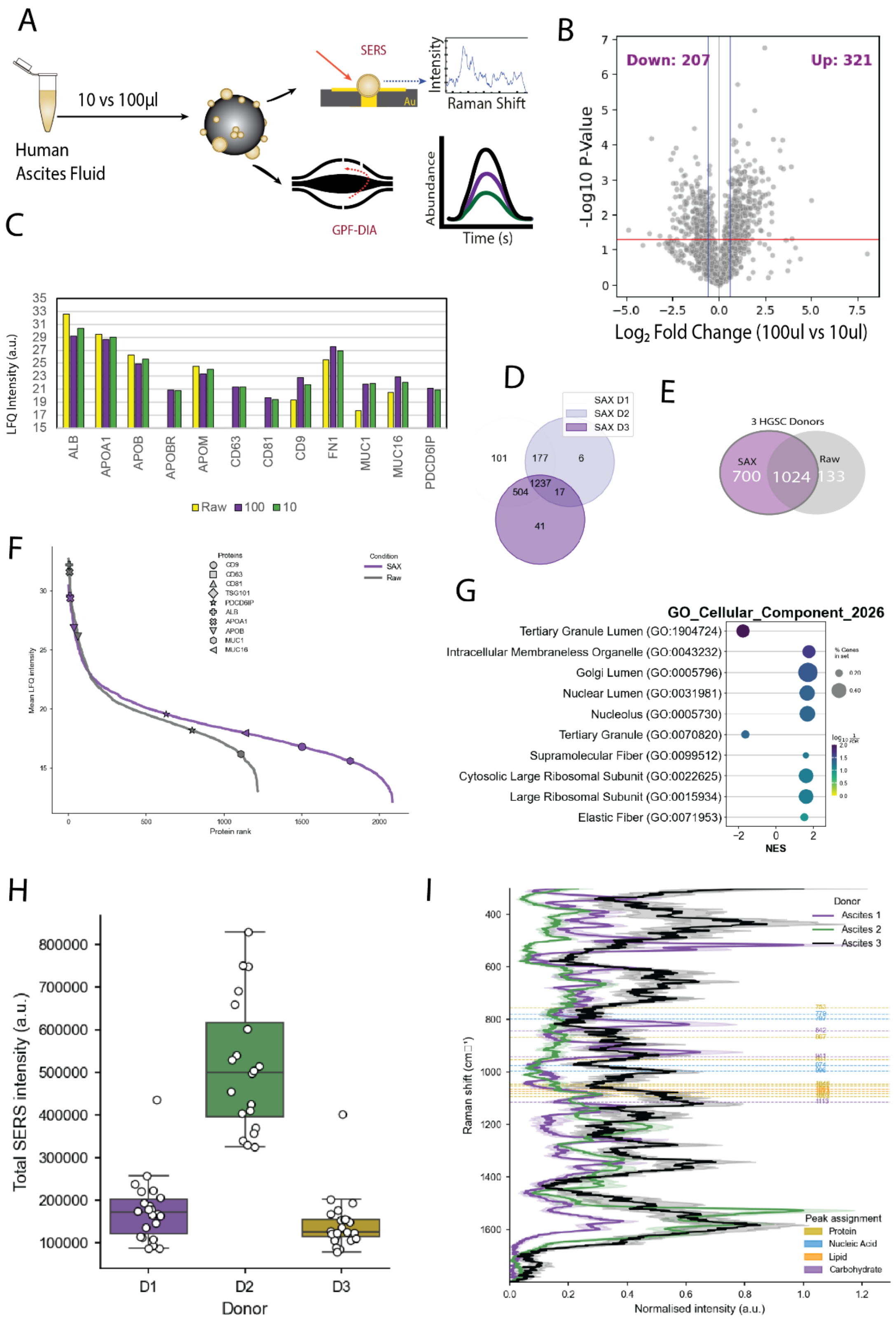

**Figure S5. Feasibility of parallel proteomic and single-EV SERS profiling from a single SAX isolation of human ascites EVs.** *To evaluate whether a single SAX-enriched EV isolation could support parallel molecular profiling, human ascites EVs were isolated from 10 or 100 µL input volumes and split for gas-phase fractionation DIA proteomics and single-EV SERS.* ***(A)*** *Workflow schematic illustrating SAX-based EV enrichment from human ascites fluid followed by parallel GPF-DIA proteomics and single-EV SERS analysis.* ***(B)*** *Differential abundance analysis comparing EV proteomes generated from 100 µL versus 10 µL ascites input, identifying 321 proteins increased and 207 proteins decreased in the 100 µL condition.* ***(C)*** *Mean LFQ intensity values of representative contaminant-, lipoprotein-, and EV-associated proteins across raw ascites and SAX-enriched EVs generated from 100 µL or 10 µL input.* ***(D)*** *Venn analysis comparing SAX-enriched EV proteomes across three independent donors.* ***(E)*** *Venn analysis comparing proteins identified in raw ascites and SAX-enriched EVs from three independent high-grade serous carcinoma donors, showing 1,024 proteins common to raw and SAX-enriched fractions, 700 proteins uniquely identified in SAX-enriched EVs, and 133 proteins uniquely identified in raw ascites.* ***(F)*** *Protein rank-abundance plot comparing raw ascites and SAX-enriched EV fractions, highlighting increased proteomic depth and dynamic range following SAX enrichment. Canonical EV-associated proteins and common plasma/lipoprotein-associated proteins are indicated by symbol.* ***(G)*** *GOCC enrichment analysis of proteins increased in SAX-enriched EVs relative to raw ascites.* ***(H)*** *Total SERS intensity of single EVs profiled from three independent donor ascites samples.* ***(I)*** *Representative normalized SERS spectra from SAX-enriched EVs across the three donor samples, with annotated spectral bands corresponding to major biochemical classes, including proteins, nucleic acids, lipids, and carbohydrates. Data in* ***(H)*** *are shown as box plots with median and interquartile range.*

Supplemental Figures

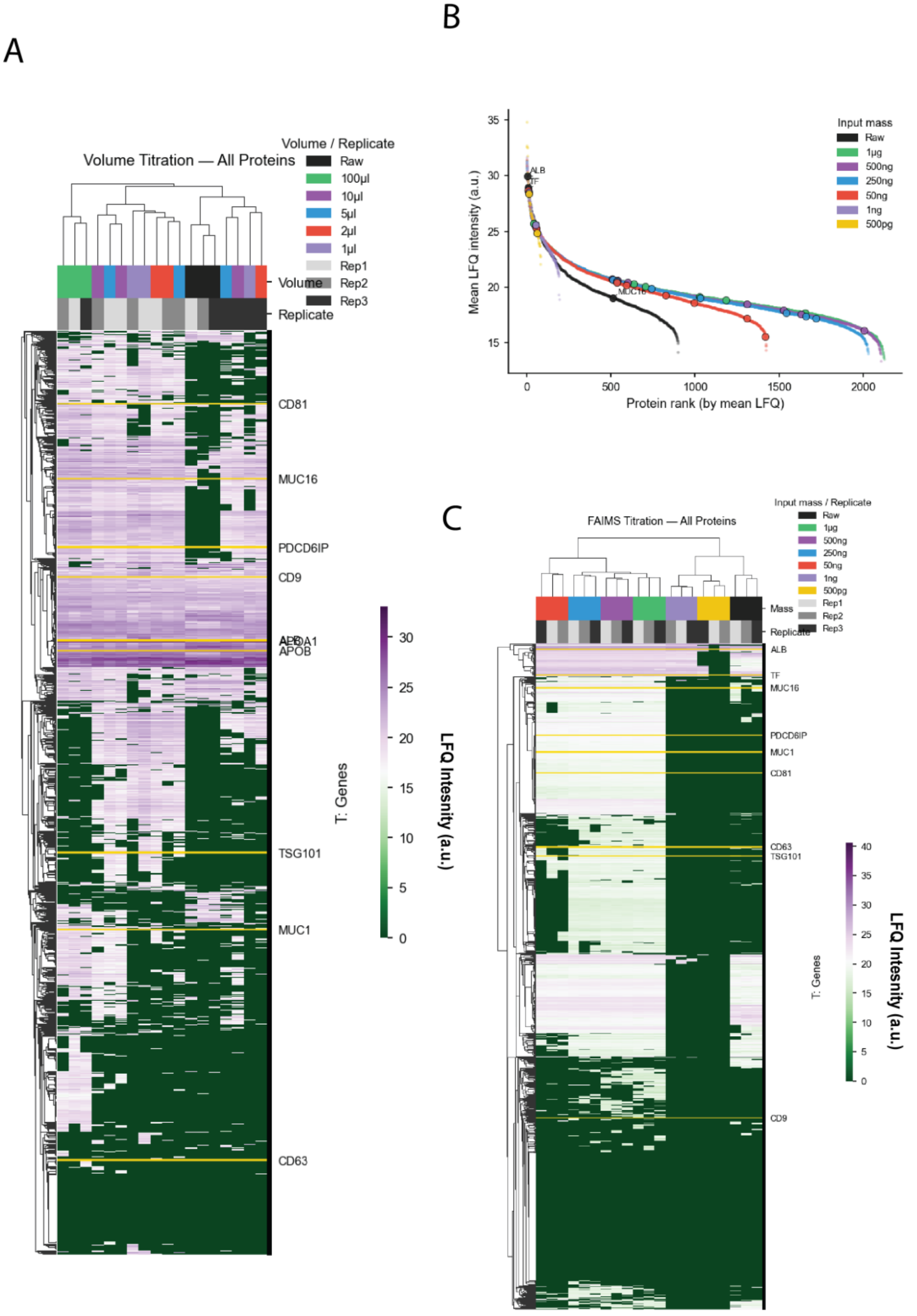

**Figure S6. Titration of ascites input volume and peptide injection mass for SAX-EV proteomic profiling.** *Additional analyses were performed to evaluate how starting ascites volume and peptide injection mass influence the depth and reproducibility of SAX-enriched EV proteomic profiling.* ***(A)*** *Unsupervised clustering and heatmap of LFQ intensities for all quantified proteins across the input-volume titration experiment, including raw ascites and SAX-enriched EVs generated from 1, 2, 5, 10, and 100 µL of human ascites fluid. Replicates are indicated in the annotation bar, and selected EV-associated, tumour-associated, apolipoprotein, and contaminant proteins are highlighted.* ***(B)*** *Protein rank-abundance analysis showing mean LFQ intensity as a function of protein rank across the peptide injection-mass titration experiment. SAX-enriched EV peptide inputs ranging from 500 pg to 1 µg are compared with raw ascites.* ***(C)*** *Unsupervised clustering and heatmap of LFQ intensities for all quantified proteins across the FAIMS-assisted injection-mass titration experiment, including raw ascites and SAX-enriched EV peptide inputs of 500 pg, 1 ng, 50 ng, 250 ng, 500 ng, and 1 µg. Selected EV-associated, tumour-associated, apolipoprotein, and contaminant proteins are highlighted.*
